# Maternal diet-induced obesity during pregnancy alters lipid supply to fetuses and changes the cardiac tissue lipidome in a sex-dependent manner

**DOI:** 10.1101/2021.04.12.439435

**Authors:** Lucas C. Pantaleão, Isabella Inzani, Samuel Furse, Elena Loche, Antonia Hufnagel, Thomas Ashmore, Heather L. Blackmore, Benjamin Jenkins, Asha A. M. Carpenter, Ania Wilczynska, Martin Bushell, Albert Koulman, Denise S. Fernandez-Twinn, Susan E. Ozanne

**Affiliations:** University of Cambridge Metabolic Research Laboratories and MRC Metabolic Diseases Unit, Level 4, Addenbrooke’s Hospital, Cambridge, Cambridgeshire, United Kingdom, CB22 0QQ.; CRUK Beatson Institute, Garscube Estate, Switchback Road, Bearsden, Glasgow, United Kingdom, G61 1BD

**Keywords:** Maternal obesity, Fetal heart, Heart metabolism, Lipidomics, Transcriptomics

## Abstract

Maternal obesity during pregnancy has immediate and long-term detrimental effects on the offspring heart. In this study, we characterized the cardiac and circulatory lipid profiles in fetuses of diet-induced obese pregnant mice and established the changes in lipid abundance and fetal cardiac transcriptomics. We used untargeted and targeted lipidomics and transcriptomics to define changes in the serum and cardiac lipid composition and fatty acid metabolism in male and female fetuses. From these analyses we observed: (1) maternal obesity affects the maternal and fetal serum lipidome distinctly; (2) female heart lipidomes are more sensitive to maternal obesity than male fetuses; (3) changes in lipid supply might contribute to early expression of lipolytic genes in mouse hearts exposed to maternal obesity. These results highlight the existence of sexually dimorphic responses of the fetal heart to the same *in utero* obesogenic environment and identify lipids species that might mediate programming of cardiovascular health.

## INTRODUCTION

Mammalian heart development and maturation involve a complex array of processes that are only completed postnatally, when increased systemic demands and changes in substrate and oxygen availability promote major cardiac remodelling (Reviewed by Piquereau and Ventura-Clapier, 2018). Appropriate and regulated flow of hormones, nutrients, metabolites and absorbed gases into fetal tissues is required to ensure that intrauterine development is achieved in an appropriate time-sensitive manner. Therefore, adverse gestational conditions – such as maternal obesity – can disrupt maternal/fetal molecule interchange, leading to impaired fetal development, which can have long-term impacts on cardio-metabolic health postnatally (Dong *et al*, 2012; Zambrano and Nathanielsz, 2013). Such a causal link between maternal metabolic status and lifelong offspring health and disease is encompassed in what has been termed the Developmental Origins of Health and Disease (Barker, 2007).

Maternal obesity during gestation is one condition that has been shown to raise the risk of non-communicable diseases in the expectant mother and her children. Numerous studies in humans and animal models suggest that obesity during pregnancy has immediate and long-term detrimental effects including increased risk of congenital heart disease (Helle and Priest, 2020), and increased susceptibility of the offspring to cardiometabolic abnormalities postnatally (Guénard *et al*., 2013, Loche *et al*, 2018). This is of particular importance, as recent data indicates that around 50% of pregnant women in developed countries are currently either overweight or obese, and cardiovascular diseases are a leading cause of death worldwide (NMPA Project Team, 2019; GBD, 2017).

Mild hypoxia, high maternal insulin and leptin levels, as well as changes in nutrient and metabolite availability have been implicated as causal factors, mediating the effects of maternal obesity on the fetus and its long-term health (Howell and Powell, 2017). However, there is limited data in relation to the molecular consequences of such exposure on the fetal heart.

Although a small number of studies have provided some evidence that maternal obesity affects the fetal cardiac transcriptome and protein profile, these studies do not explain the whole complexity of changes in the cardiac phenotype. In addition, there is very limited data on the contribution of the maternal and fetal lipidome to programming mechanisms (Catalano and Shankar, 2017).

Lipids are a complex group of structural, energy and signalling molecules involved in a variety of physiological, metabolic and pathological processes. Changes in the murine heart lipidome have been shown to initiate and promote inflammatory reactions after infarction (Halade *et al*, 2017), and cellular lipid composition has been associated with the distinction between physiological and pathological cardiac hypertrophy and with prognosis of cardiac disease (Tham *et al*, 2018, Le *et al*, 2014). Moreover, recent studies explored the association of the cardiac lipidome with life stage progression, showing how changes in intracellular lipids contribute to heart maturation at birth (Walejco *et al*, 2018) and the impact of ageing on the heart (Eum *et al*, 2020).

Despite the growing interest in lipid profiling in health and disease, the study of fetal cardiac lipids in maternal obesity models remains largely unexplored. The aim of the current study was therefore to use lipidomics and cardiac transcriptomics to identify lipid pathways that may be associated with developmental phenotypes that lead to chronic diseases later in life. In addition, given the growing evidence for sex-specific differences in the programming field, a further aim was to establish if any of the responses were sexually dimorphic. As circulating lipids are often used as biomarkers of cardiovascular health, we also sought to investigate associations between maternal and fetal serum lipidomes.

## RESULTS

### Maternal obesity affects female, but not male, mouse offspring heart morphology

A well-established diet-induced maternal obesity model in which the female mouse develops obesity and gestational diabetes was used to investigate the effects of maternal obesity on the offspring heart. At gestational day 18.5, male and female fetuses from obese dams were smaller than controls (Figure 1A), and although their heart weights were not significantly different from control hearts in either absolute or relative terms (Figure 1B, 1C), obese female, but not male, fetal cardiomyocytes were smaller, as observed by the increased number of nuclei detected per histological section area (Figure 1D).

**Figure 1.**
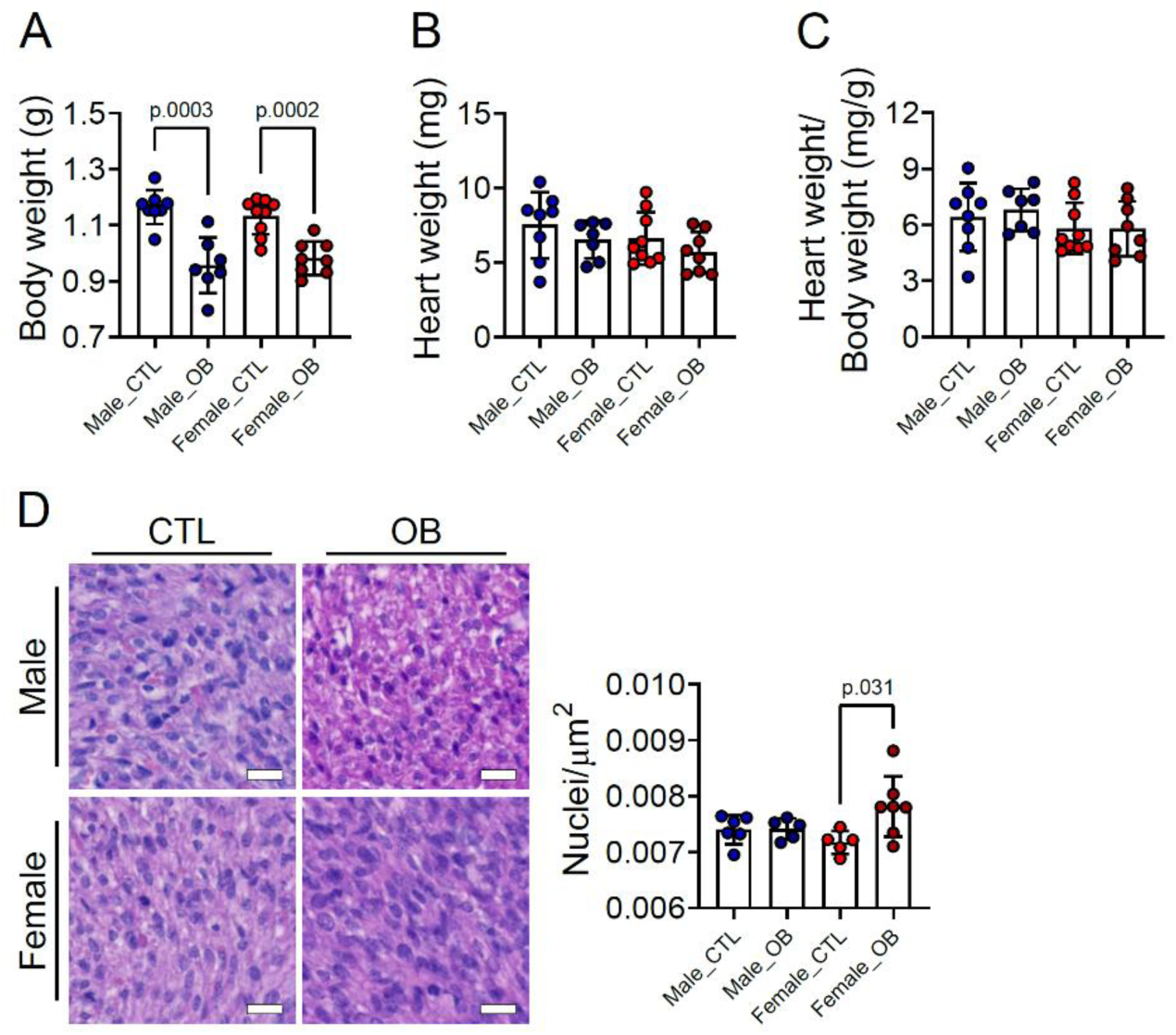
Fetal characteristics at gestational day 18.5. (A) Body weight of male and female fetuses from healthy control (CTL) and obese (OB) mouse dams at gestational day 18.5. (B-C) Heart weight and heart weight/body weight ratio of male and female fetuses from CTL and OB dams at gestational day 18.5. Male CTL n=8, male OB n=7, female CTL n=9, female OB n=8. (D) Representative images and quantification of cell nuclei count per µm^2^. Histological sections stained with haematoxylin and eosin of male and female fetuses from CTL and OB dams at gestational day 18.5. Male CTL n=6, male OB n=6, female CTL n=5, female OB n=7. Scale bar indicates 20 µm. p-value calculated by Student t-test.

### Maternal obesity drives changes to the lipid composition of maternal and fetal serum

Maternal and fetal serum lipidomes were obtained by direct infusion high-resolution mass spectrometry, a rapid method used to profile the lipids in an organic extract. Full data and annotation of isobaric signals are available in the supplementary information (Supplementary file 1, Figure 2–source Data 1). We used Principal Component Analysis (PCA) to identify orthogonal distance and relatedness amongst individual fetal and maternal serum lipidomes.

This multivariate analysis demonstrated that there was a clear distinction between maternal and fetal serum lipid profiles, regardless of the maternal nutritional status or offspring sex (Figure 2–figure supplement 1A). In order to test the hypothesis that the lipid composition in the serum of the dams and her male and female fetuses differed between control and obese mothers, PCAs of just these pairs of groups were performed. These suggested that there was clear segregation in each case driven by maternal dietary status (Figure 2–figure supplement 1B, C, D).

**Figure 2–figure supplement 1.**
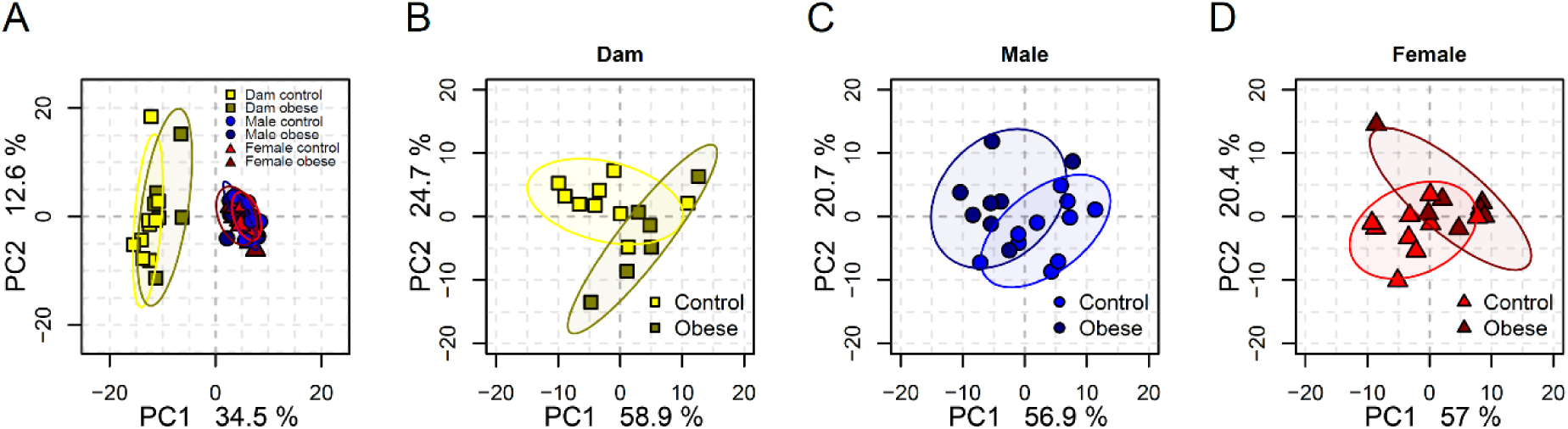
(A) PCA plots showing the PC1 and PC2 scores for individual dam and fetal serum lipidomes at gestational day 18.5. (B-D) PCA plots showing the PC1 and PC2 scores for individual dam (B), male (C) and female fetal (D) serum lipidomes.

In order to identify the lipid pathways altered, we summed the abundance of the lipid variables in each lipid class (head group, assuming even chain length for the fatty acid, see supplementary files 1 and 2 for lipid signals used for each class) and calculated which classes differed in abundance according to maternal status. This showed that cholesteryl esters, ceramides, and sphingomyelins were more abundant in serum from obese dams than in serum from controls (Figure 2A). Amongst fetal serum, we observed changes to the abundance of lipid classes that were generally similar between males and females, though different classes reached statistical significance in males (triglycerides, phosphatidic acids and the ratio of triglycerides to phospholipids) (Figure 2B) and females (phosphatidylcholines/phosphatidylethanolamines, phosphatidic acids and ceramides) (Figure 2C). We then used factorial analysis to show that the abundance of triglycerides, phosphatidylcholines/phosphatidylethanolamines and ceramides were all significantly regulated by maternal diet (Figure 2D). We also observed sex differences in cholesteryl esters, with males showing higher relative abundance compared to females (Figure 2D, see also Figure 2–figure supplement 2A).

**Figure 2.**
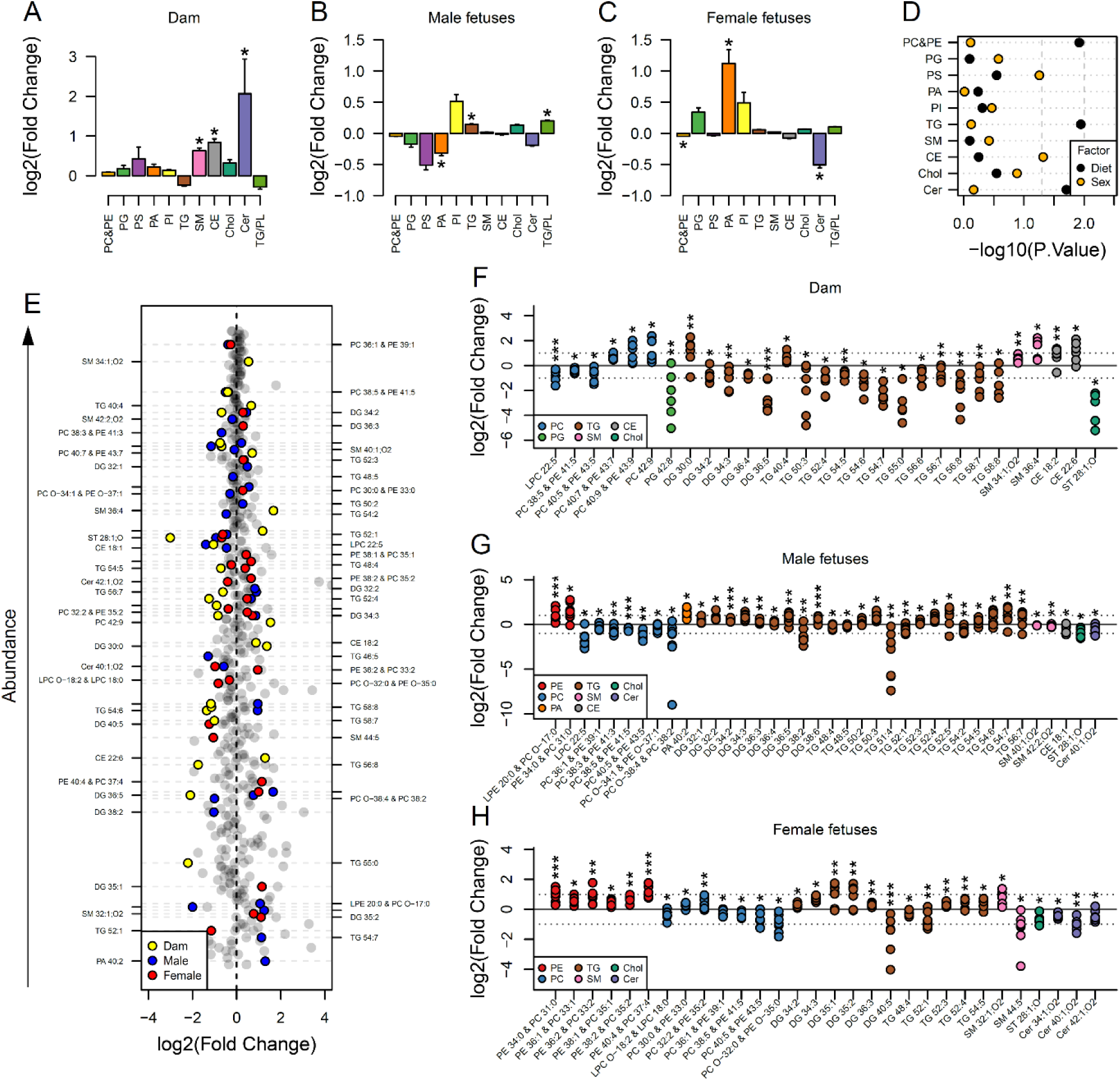
Maternal and fetal serum lipidome measured by direct infusion mass spectrometry. (A-C) Relative changes in serum lipid classes abundance in obese dams (A), male (B) and female (C) obese fetuses. Values are mean + SE. *p<0.05 calculated by Student t-test or Mann-Whitney test. (D) Influence of maternal diet and sex on fetal serum lipid classes abundance as calculated by factorial ANOVA. (E) Regulation of maternal and fetal serum lipid species ranked according to their abundance. Coloured dots represent statistically regulated species as calculated by univariate Student t-test (p<0.05) and PLS-DA VIP (vip score>1) in maternal or fetal OB serum compared to CTL. (F-H) Serum levels of regulated lipids from obese dams (F) and from male (G) and female (H) fetuses of obese dams at gestational day 18.5. Each dot represents a result from one obese heart, relative to the average of results for individual lipids in the control group (straight line). Dam CTL n=9, dam OB n=6, male fetuses CTL n=10, male fetuses OB n = 8, female fetuses CTL n=10, female fetuses OB n=7; * p<0.05, ** p<0.01, *** p<0.001 calculated by Student t-test. In figures A-D: PE, phosphatidylethanolamines/odd chain phosphatidylcholines; PC, phosphatidylcholines/odd-chain phosphatidylethanolamines; PG, phosphatidylglycerols; PS, phosphatidylserines; PA, phosphatidic acids; PI, phosphatidylinositols; TG, monoglycerides, diglycerides and triglycerides; SM, sphingomyelins; CE, cholesteryl esters; Cer, ceramides; PL, phospholipids. In figures E-H, other isobaric lipids can contribute to these signals (Supplementary file 1). See also Figure 2–figure supplement 1 and Figure 2–figure supplement 2. Full data is available in Figure 2–source Data 1.

### Specific lipid profiles differ between maternal and fetal serum

Lipid classes consist of several lipid isoforms that comprise different fatty acid residues. To assess the differences in lipids at the individual level, we used both univariate and multivariate models to identify differences between serum from obese and control dams and their fetuses. Although ceramides were more abundant at the class level in obese dam serum, no ceramide isoform was statistically significantly different between groups (Figure 2E, 2F). At the individual isoform level, campesterol (ST 28:1;O) and many unsaturated triglycerides were less abundant. Similar to changes observed at the class level, the abundance of two isoforms of sphingomyelin, and several individual cholesteryl esters, was also increased in the sera of obese mothers. Some phosphatidylcholines/odd chain phosphatidylethanolamines were more abundant, although two – PC 40:5/PE 43:5 and PC 38:5/PE 41:5 – and *lyso*-phosphatidylcholine (LPC) 22:5 were less abundant.

We then explored if a subset of lipids was transported from the maternal to the fetal circulation using linear regression. We showed that only a few maternal phospholipids and *lyso*-phospholipids species – comprising LPC 18:2, 20:4, 22:5 and 22:6 – and campesterol were significantly correlated with the same species in both male and female fetal serum (Figure 2–figure supplement 2B, 2C). Contrasting to the signature observed in dam serum, naturally highly abundant triglyceride isoforms had increased abundance in both male and female fetuses with only a handful of exceptions (e.g., TG 54:2 and TG 48:5 in males). Phosphatidylethanolamine/odd chain phosphatidylcholines isoforms were also more abundant in both males and females, with females showing increased abundance of the greatest number of species, and PE 34:0/PC 31:0 being regulated in a sex-independent manner (Figure 2E, 2G, 2H). Odd chain fatty acid containing phosphatidylcholines are isobaric with phosphatidylethanolamines (see supplementary file 2), however no other evidence from other lipid classes suggested a change in odd chain fatty acid metabolism. Consistent with the dam serum data, several phosphatidylcholine/odd chain phosphatidylethanolamines isoforms were less abundant in male and female fetuses.

**Figure 2–figure supplement 2.**
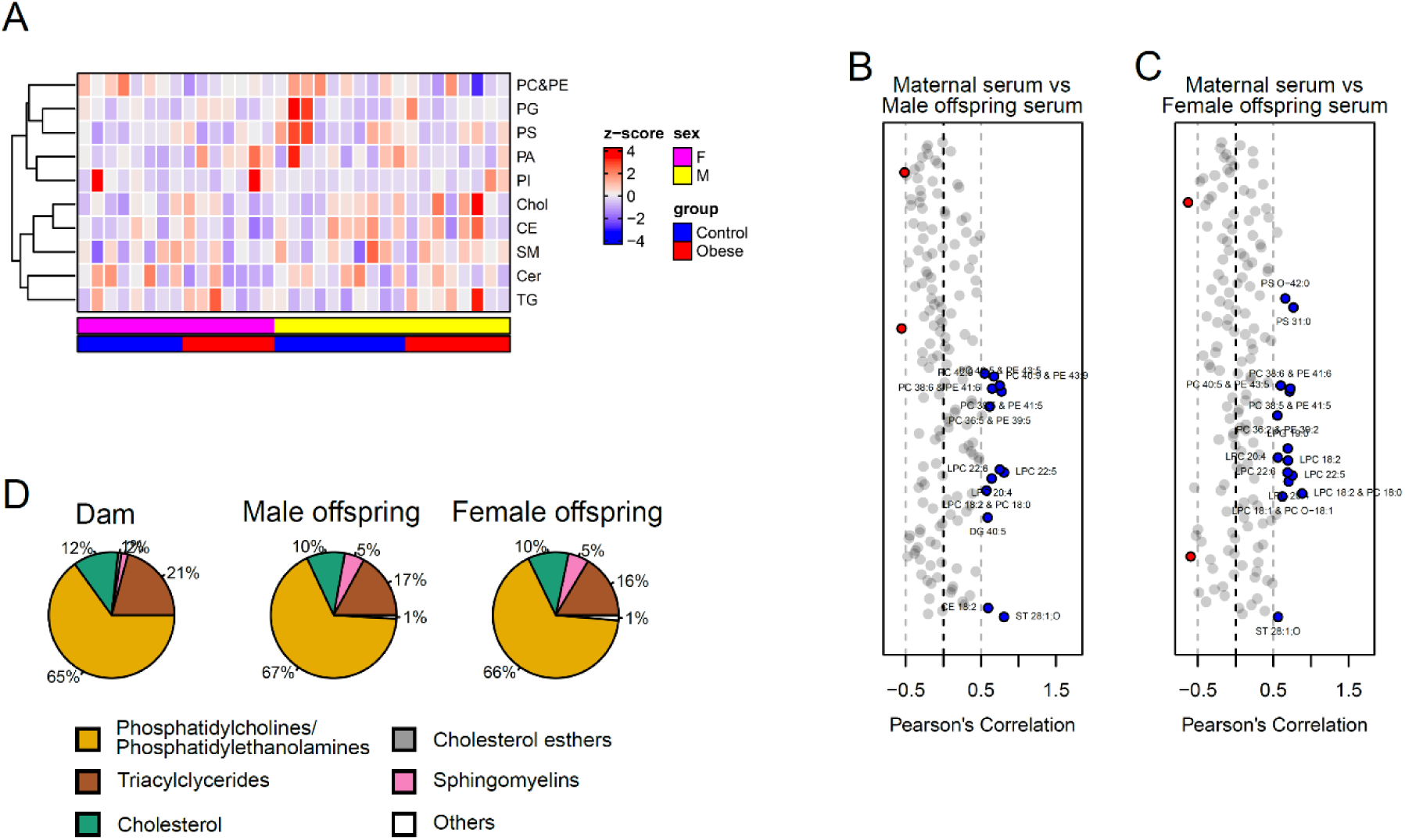
(A) Heatmap showing lipid classes serum levels in male and female E18.5 fetuses. (B-C) Pearson’s correlation between individual lipid species in maternal serum and male (B) and female (C) fetal serum at gestational day 18.5. Blue dots represent positively correlated lipid species between maternal and fetal serum deemed statistically significant (p<0.05). Red dots represent negatively correlated lipid species between maternal and fetal serum deemed statistically significant (p<0.05). (D) Relative amount of different lipid classes in maternal and fetal serum.

### Maternal obesity is associated with the fatty acid composition of phospholipids in maternal and fetal serum

The bulk of the total of serum lipid species detected were glycerides, phospholipids and cholesterol (Figure 2–figure supplement 2D), with glycerides, phospholipids and *lyso*-phospholipids comprising three and two fatty acids respectively, that are covalently bound to glycerol. As we observed differences in the abundance of individual lipid isoforms in maternal and fetal serum, we sought to elucidate whether the distinct signatures observed in fetuses from obese dams would also translate into an imbalance in the distribution of fatty acid residues in phospholipids. When clustered according to the number of double bonds as either saturated, monounsaturated or polyunsaturated residues, we saw that the relative abundances of these were not significantly affected by maternal diet either in maternal or fetal serum (Figure 3A). We also noted that fatty acids from phospholipids with a chain length shorter than 18 carbons were generally more abundant in obese serum, whereas the abundance of longer molecules tended to be lower (Figure 3B).

**Figure 3.**
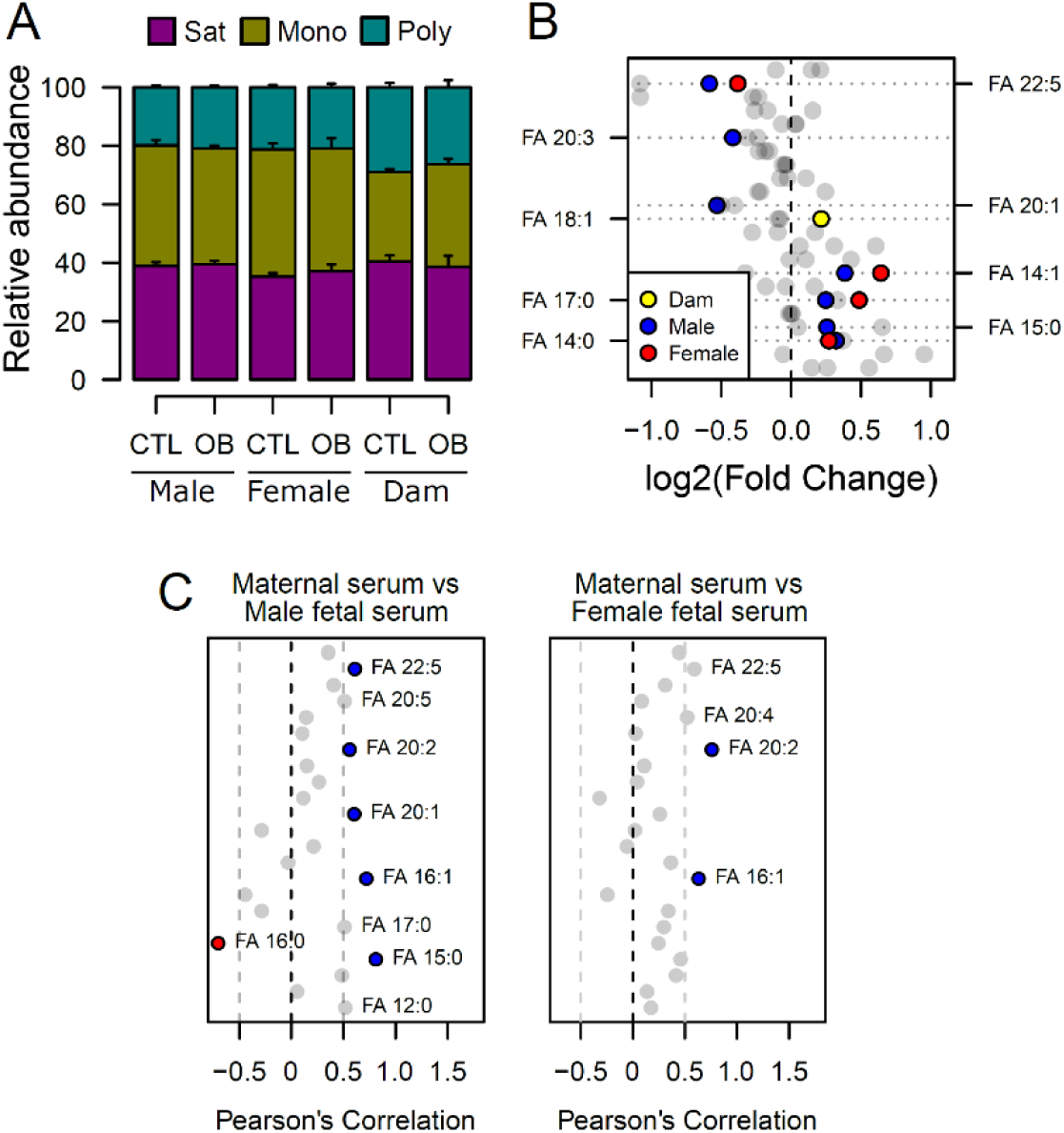
Fatty acid composition of serum phospholipids measured by direct infusion mass spectrometry using in-source CID fragmentation. (A) Grouped saturated, monounsaturated and polyunsaturated fatty acids content in maternal, male and female fetal serum at gestational day 18.5. Values are mean + SE. (B) Regulation of maternal and fetal serum fatty acids. Coloured dots represent statistically regulated fatty acids as calculated by univariate Student t-test or Mann-Whitney test (p<0.05) in maternal or fetal OB serum compared to CTL. (C) Pearson’s correlation between maternal serum fatty acids and the same fatty acids detected in the fetal serum. Blue and red dots represent species with significant positive and negative association (p<0.05). Dam CTL n=8, dam OB n=6, male fetuses CTL n=10, male fetuses OB n = 8, female fetuses CTL n=8, female fetuses OB n=6. See also Figure 3–figure supplement 1 and Figure 3–figure supplement 2. Full data is available in Figure 3–source Data 1

At the individual level, we again identified a distinction between maternal and fetal serum profiles, with phospholipid fatty acid residues being differentially regulated (Figure 3B, see also Figure 3–figure supplement 1).

**Figure 3–figure supplement 1.**
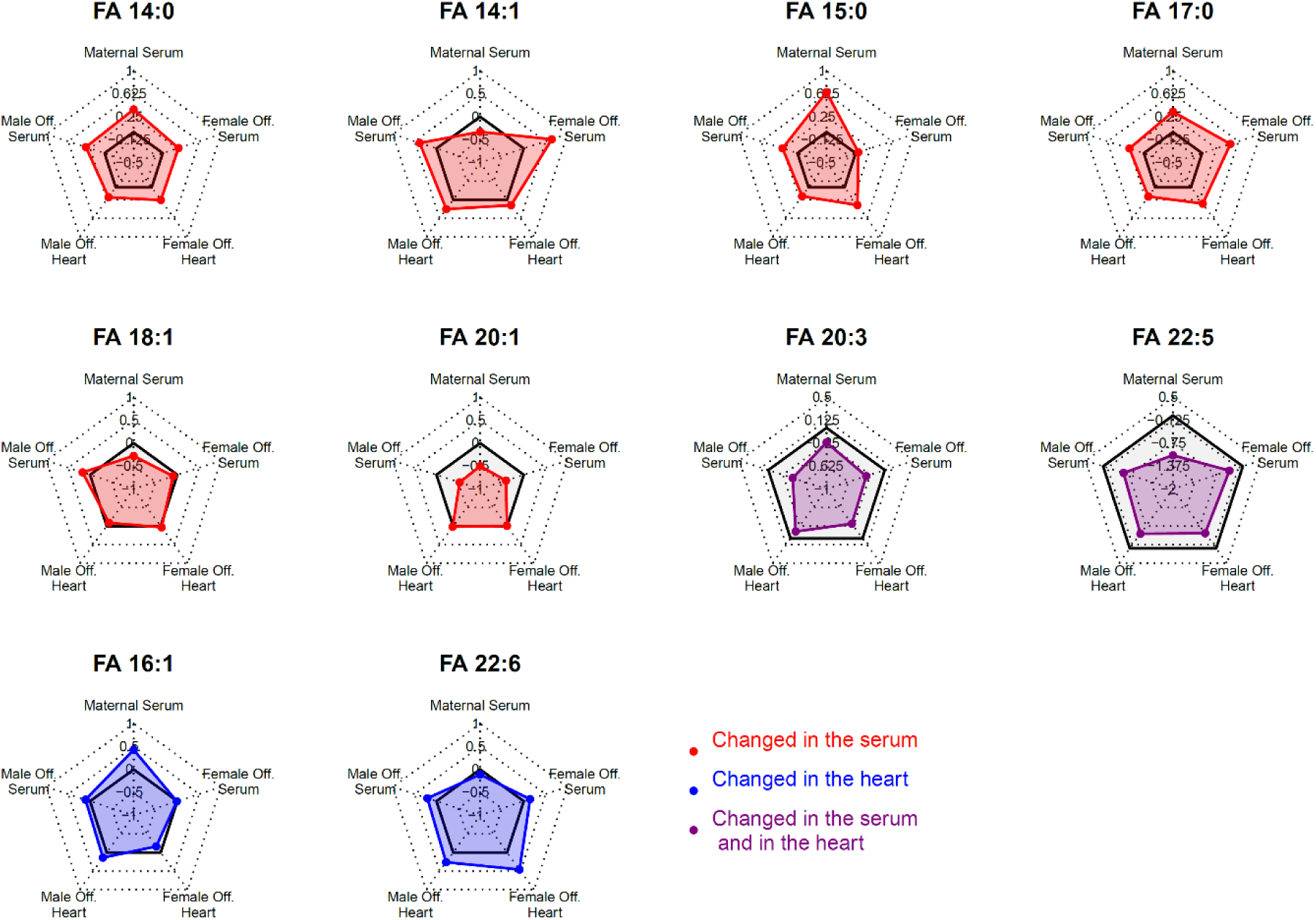
Radar plots showing log2 of fold change of fatty acids statistically changed in the serum or in the heart of fetuses from obese dams in different compartments. Grey shaded area indicates log2 fold change smaller than 0.

In line with the lower polyunsaturated phospholipid levels, those containing n-3 docosapentaenoic acid (DPA) (22:5) were less abundant, and those with saturated fatty acids myristic (14:0) and margaric (17:0), and monounsaturated myristoleic acid (14:1) were more abundant in the fetal sera of both sexes in response to maternal obesity. In contrast, oleic acid (18:1) from phospholipids was more abundant in the serum of obese dams only. Despite the differences observed, the fetal levels of a few residues were significantly correlated with the maternal ones, and an overall trend for positive correlation between fatty acids levels in maternal and fetal sera was observed (Figure 3C). By generating correlation matrices with group independent variables using Euclidian clustering, we also observed that several saturated and monounsaturated fatty acid residues were positively correlated and tended to cluster together (Figure 3–figure supplement 2). Similarly, polyunsaturated fatty acids established clusters and were positively correlated. Several saturated and monounsaturated fatty acids were negatively correlated to polyunsaturated species.

**Figure 3–figure supplement 2.**
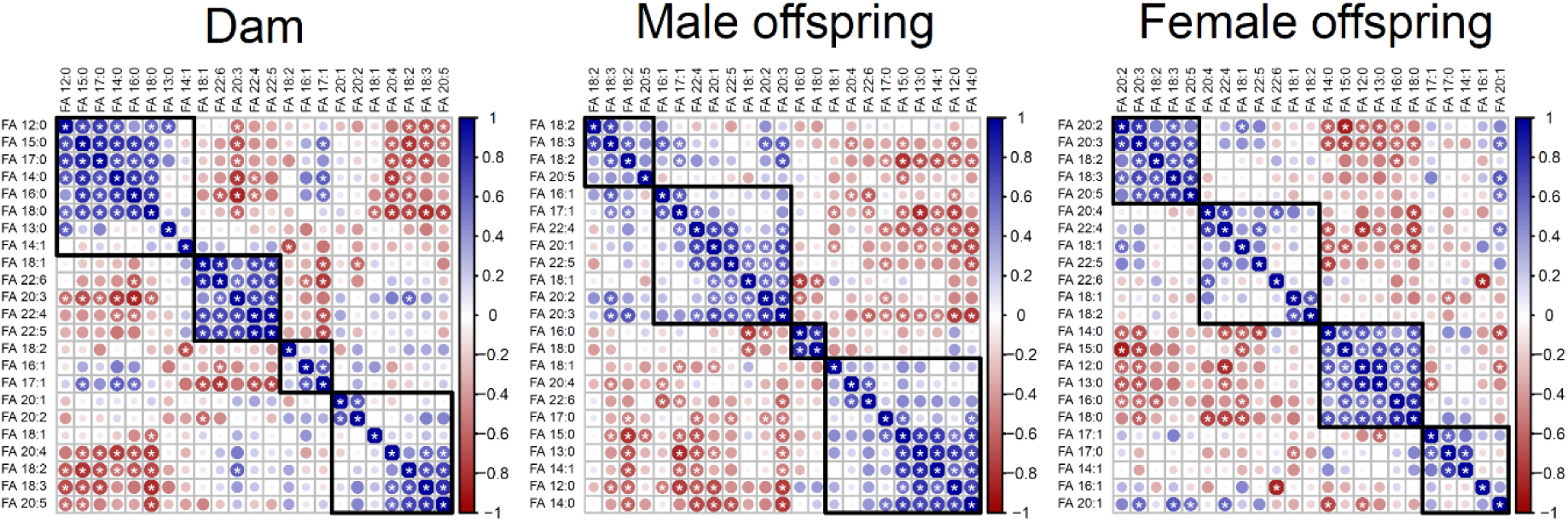
Correlation matrices showing Pearson’s correlation between cardiac fatty acids in dams and fetuses. Fatty acids grouped following Euclidian clusterization.

### Maternal obesity sex-specifically affects the fetal heart lipidome

We next sought to determine whether the lipid composition of fetal hearts was influenced by maternal obesity and whether its signature would follow a similar pattern to that observed in the fetal serum. Full cardiac lipidome data and annotation of isobaric signals are available in the supplementary information (Supplementary file 2, Figure 4–source Data 1 and Figure 4–source Data 2). We observed that total cholesteryl esters were less abundant in both male and female fetal hearts from obese pregnancies (Figure 4A, 4B). Total sphingomyelins were more abundant in both males and females, but the difference was only statistically significant in female hearts (Figure 4B). The observation that fewer lipid classes were perturbed in males than in females in response to maternal obesity was reproduced at the individual species level. Through PCAs, we observed a weaker degree of separation between control and obese individual lipidomes in males (Figure 4C) when compared to females (Figure 4D). In contrast, female heart lipidomes showed clear distinction between control and obese hearts, with more lipid species being significantly different between groups (42 compared to 18 in males) (Figures 4E, 4F and 4G).

**Figure 4.**
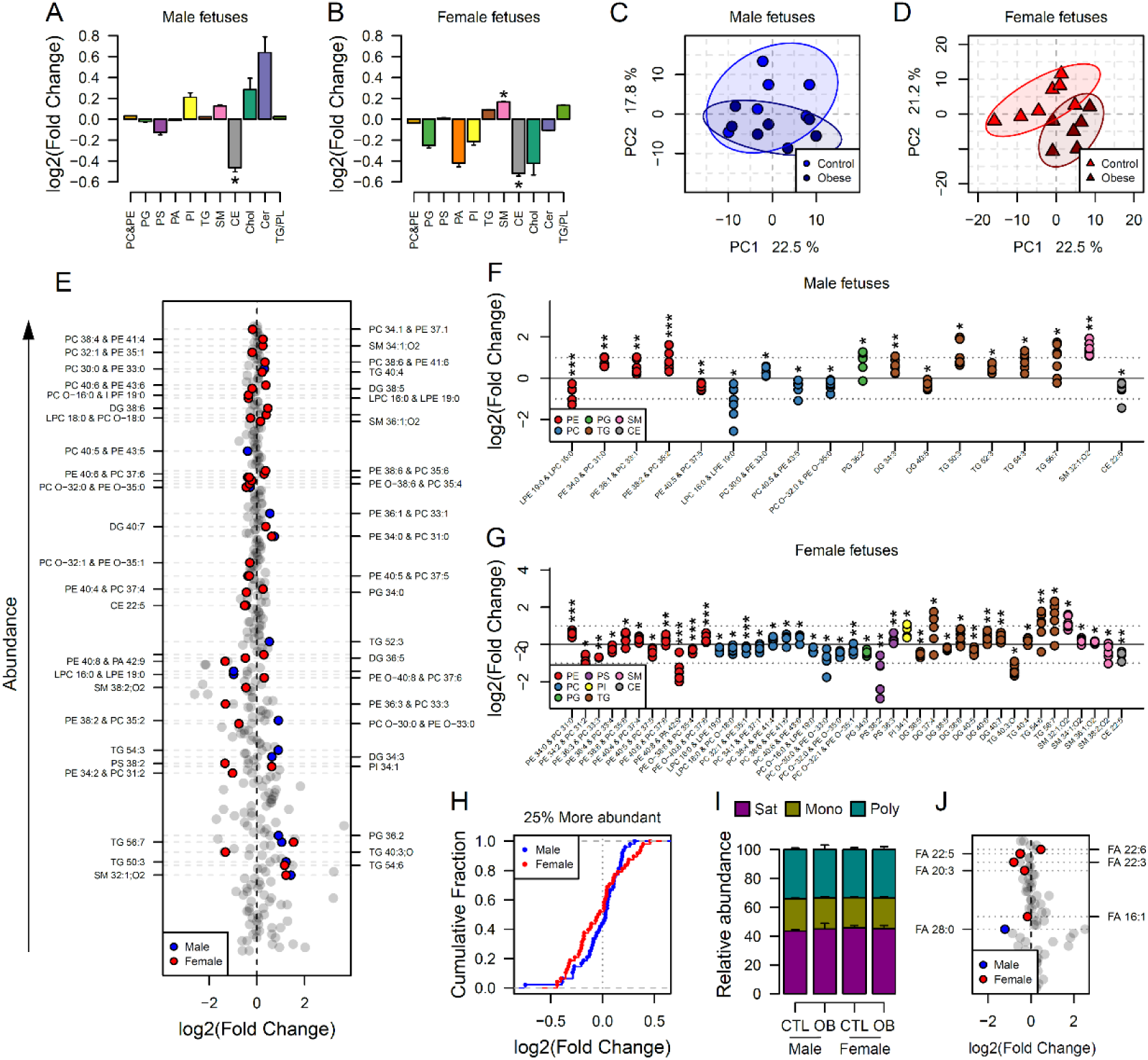
Maternal and fetal cardiac lipidome. (A-B) Relative changes in cardiac lipid classes in male (A) and female (B) fetuses from obese dams. Values are mean + SE. *p<0.05 calculated by Student t-test or Mann-Whitney test. (C-D) PCA plots showing the PC1 and PC2 scores for individual male (C) and female (D) cardiac lipidomes. (E) Regulation of fetal cardiac lipid species ranked according to their abundance. Coloured dots represent statistically regulated species as calculated by univariate Student t-test (p<0.05) and PLS-DA VIP (vip score>1) in fetal OB hearts compared to CTL. (F-G) Cardiac levels of regulated lipids from male (F) and female (G) fetuses of obese dams at gestational day 18.5. Each dot represents a result from one obese heart, relative to the average of results for individual lipids in the control group (straight line). Male fetuses CTL n=6, male fetuses OB n=7, female fetuses CTL n=7, female fetuses OB n=6. * p<0.05, ** p<0.01, *** p<0.001 calculated by Student t-test. (H) Cumulative frequency of cardiac lipid species according to the log2 of the fold change in abundance between male and female fetuses from obese and control dams. (I) Grouped saturated, monounsaturated and polyunsaturated fatty acids content in male and female fetal hearts at gestational day 18.5. (J) Regulation of maternal and fetal serum fatty acids. Coloured dots represent statistically regulated fatty acids as calculated by univariate Student t-test or Mann-Whitney test (p<0.05) in fetal OB hearts compared to CTL. Male fetuses CTL n=8, male fetuses OB n=6, female fetuses CTL n=7, female fetuses OB n=7. In figures A-B: PE, phosphatidylethanolamines/odd chain phosphatidylcholines; PC, phosphatidylcholines/odd-chain phosphatidylethanolamines; PC, phosphatidylcholines; PG, phosphatidylglycerols; PS, phosphatidylserines; PA, phosphatidic acids; PI, phosphatidylinositols; TG, monoglycerides, diglycerides and triglycerides; SM, sphingomyelins; CE, cholesteryl esters; Cer, ceramides; PL, phospholipids. In figures E-G, other isobaric lipids can contribute to these signals (Supplementary file 2). See also Figure 4–figure supplement 1 and Figure 4–figure supplement 2. Full data is available in Figure 4–source Data 1 and Figure 4–source Data 2.

Looking at the most abundant cardiac lipids, we observed that female hearts were more sensitive to change as a consequence of maternal obesity, with most lipids having greater fold change compared to male hearts (Figure 4H, see also Figure 3–figure supplement 1 and Figure 4–figure supplement 1).

**Figure 4–figure supplement 1.**
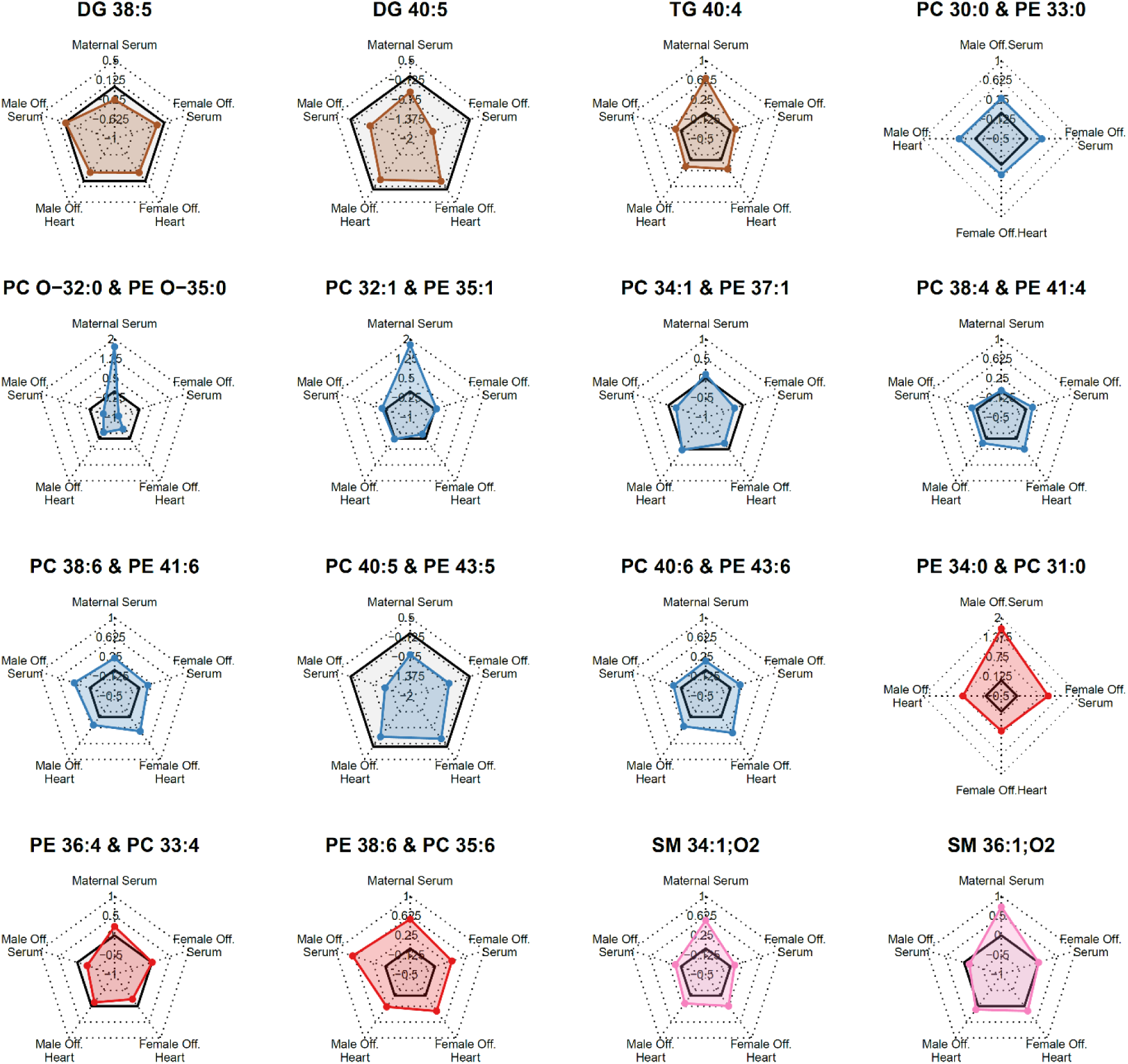
Radar plots showing log2 of fold change of most abundant statistically regulated lipids in the heart of fetuses from obese pregnancies in different compartments. Grey shaded area indicates log2 fold change smaller than 0.

Lipid ontology analysis of the fetal heart lipidome using LION (Moleenar et al., 2019) also identified more biological features significantly enriched in female hearts (Figure 4–figure supplement 2). This analysis also revealed an overall decrease in phospholipids and *lyso-*lipids and, although not quantitatively significant, an increase in triglycerides in both males and females.

**Figure 4–figure supplement 2.**
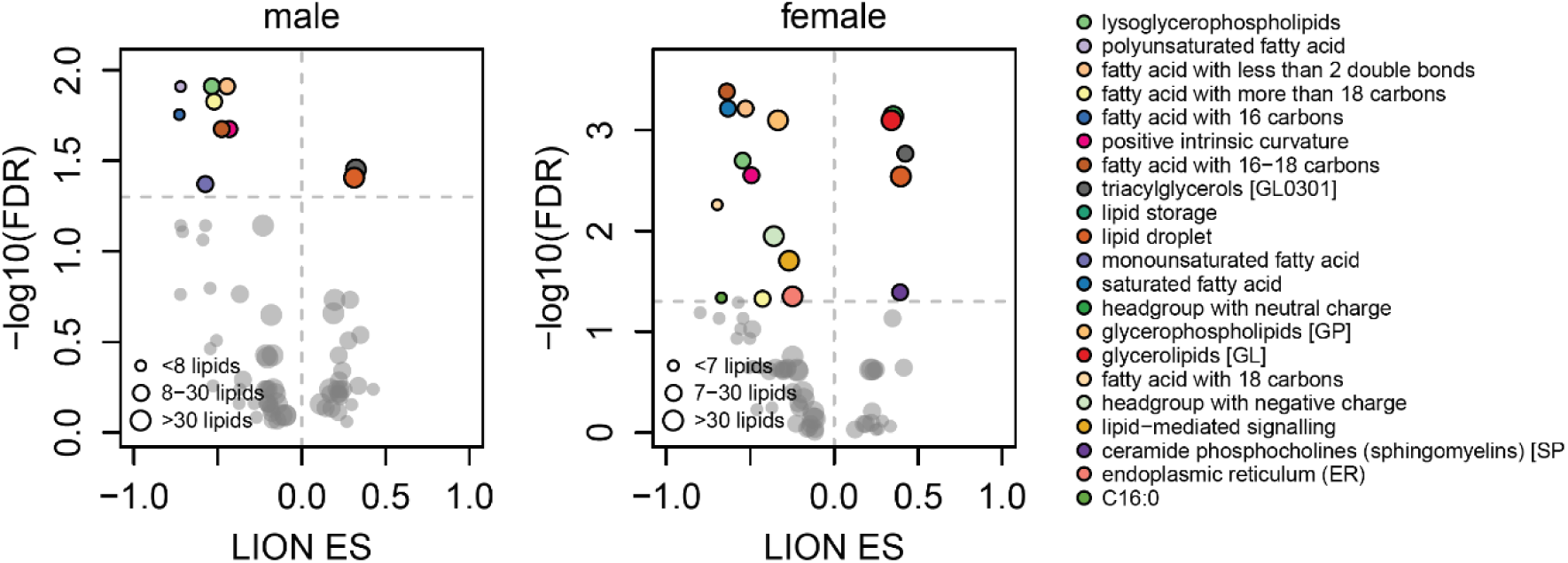
Scatterplots showing enrichment score (ES) and statistical significance of lipid ontology pathways from LION.

Consistently, most modelled triglycerides were more abundant, and most phosphatidylcholines/odd chain phosphatidylethanolamines were less abundant in both male and female hearts exposed to *in utero* obesity (Figure 4F, 4G). Phosphatidylethanolamines/odd chain phosphatidylcholines were also regulated in both male and female hearts, and sphingomyelins were regulated in females only. Regarding the fatty acid composition of phospholipids, we did not observe changes in the overall content of fatty acid groups (Figure 4I). However, at the individual level, we observed lower abundance of DPA in both male and female hearts exposed to *in utero* obesity (Figure 4J). Male offspring also exhibited lower cardiac levels of m/z=423.421, tentatively identified as the very long-chain saturated octacosanoic acid (28:0), although other metabolites can contribute to this signal. In females, we observed increased abundance of highly abundant docosahexaenoic acid (22:6) and lower abundance of palmitoleic (16:1) and fatty acids 22:3 and 20:3.

### Maternal obesity induces changes in the fetal heart transcriptome to promote lipid metabolism

Changes in the abundance of individual lipid isoforms and the fatty acid composition of lipid classes in cardiac cells could indicate lipid metabolism and cell morphology remodelling in cardiomyocytes. This led us to the hypothesis that maternal obesity caused changes in fetal heart lipid metabolism and biosynthesis. To identify the main gene pathways affected, we conducted RNA-Seq of male fetal hearts, followed by deep pathway enrichment analysis. Ingenuity Pathway Analysis revealed that a set of confidently top-regulated genes (FDR < 0.1) were associated with sterol, fatty acid and carnitine metabolism (Figure 5A), in a scenario where PPAR-alpha and HIF1A are main activated upstream transcriptional regulators, signalling by *lyso-*phosphatidylcholine (LPC) abundance is reduced and SREBP activity is downregulated (Figure 5B). Figure 5C shows expression of genes regulated by PPAR-alpha transcriptional activity, and expression of genes mapped to the main predicted IPA pathways. We later validated the expression of key genes associated with lipid metabolism in the hearts of both male and female offspring from a completely new cohort using qPCR (Figure 5D).

**Figure 5.**
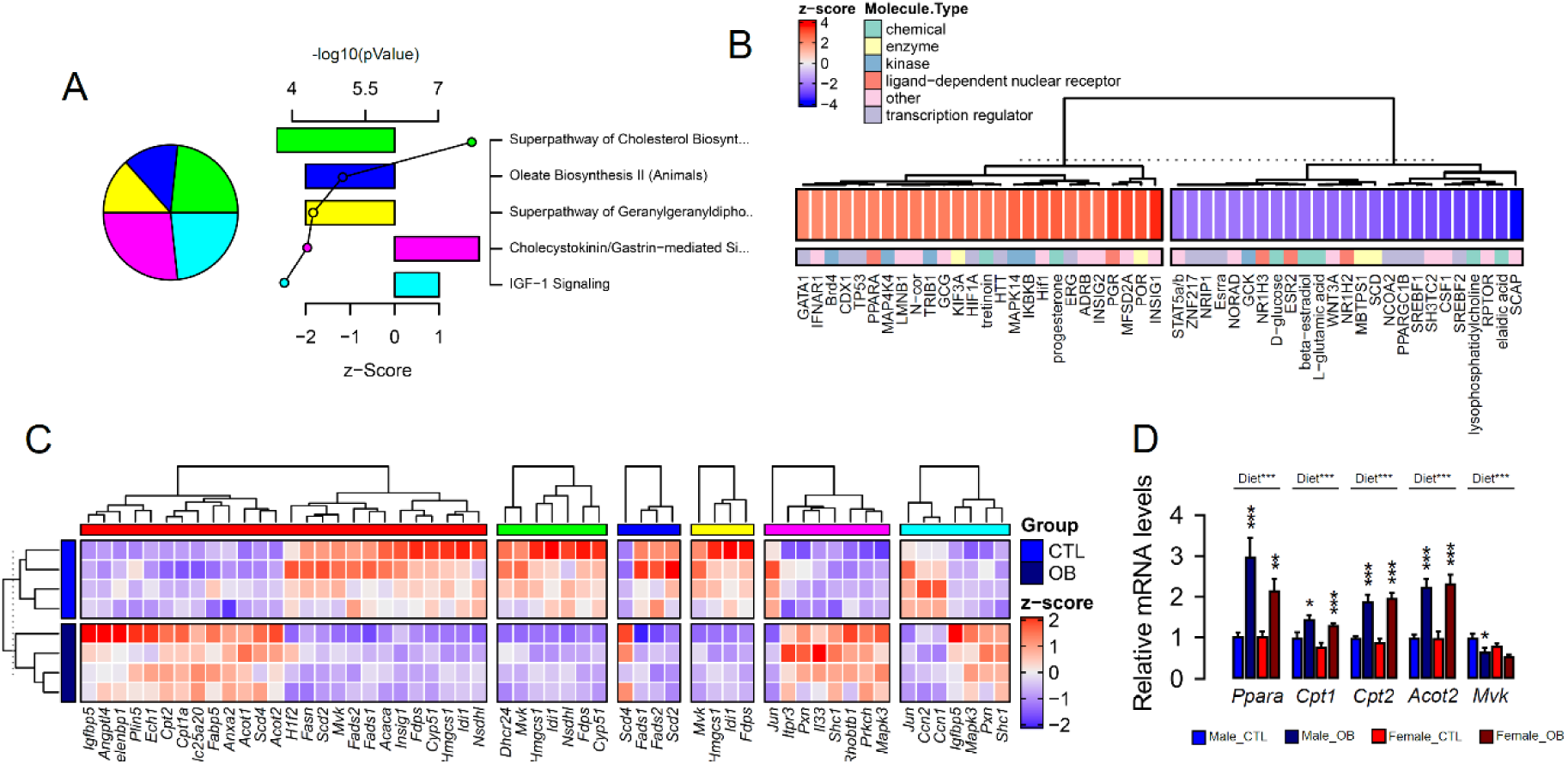
Fetal cardiac transcriptomics. (A) Top 5 regulated Ingenuity Canonical Pathways predicted by analysis of cardiac transcriptome from male fetuses from obese dams compared to fetuses from control dams. A p-value cut-off of 0.01 calculated by likelihood-ratio test was used to select regulated genes included in the IPA analysis. Pie chart represents number of genes per pathway; bars represent activation z-score per pathway; points represent p-value of enriched pathways estimated by IPA algorithm. (B) Activation z-score of top Ingenuity Upstream Regulators predicted by analysis of cardiac transcriptome from male fetuses from obese dams compared to fetuses from control dams. (C) Heatmap showing mRNA levels of genes regulated by PPAR-alpha activity (red bar), and genes mapped to “Superpathway of Cholesterol Biosynthesis” (green bar), “Oleate Biosynthesis II” (blue bar), “Superpathway of Geranylgeranyldiphosphate Biosynthesis I” (yellow bar), “Cholecystokinin/Gastrin-mediated Signalling” (pink bar) and “IGF-1 Signalling” (light blue bar) Ingenuity Canonical Pathways in male E18.5 hearts as analysed by RNA Seq. CTL n=4 and OB n=4. (D) mRNA levels of selected markers of lipid metabolism in male and female fetal heats. Male CTL n=8, male OB n=8, female CTL n=6, female OB n=11. *p<0.05, **p<0.01, ***p<0.001 by Student t-test. Diet***p<0.001 by factorial ANOVA.

### The abundance of acyl-carnitines in fetal hearts is associated with maternal obesity

Having observed changes in transcriptional activity indicating increased lipid metabolism in fetal hearts in response to maternal obesity, we conducted a final experiment to investigate whether acyl-carnitine species were also affected by maternal obesity. These comprise fatty acid residues produced during beta-oxidation and are markers of mitochondrial and peroxisomal lipid metabolism. Regardless of the lack of significant differences between sex-matched obese and control offspring (Figure 6A), we observed increased levels of total hydroxylated acyl-carnitines in response to maternal obesity by factorial ANOVA (Figure 6B). At the individual level, we found the hydroxylated acyl-carnitine C05-OH to be more abundant in both males and females (Figure 6C, 6D, 6E), and C16-OH to be less abundant in females only (Figure 6E). C12:0 and C12:1 were also more abundant, whereas C20:0 and C22:5 were less abundant in obese female hearts (Figure 6E). In males, C3:0 was less abundant, and C11:0 and C15:0 were both more abundant (Figure 6D).

**Figure 6.**
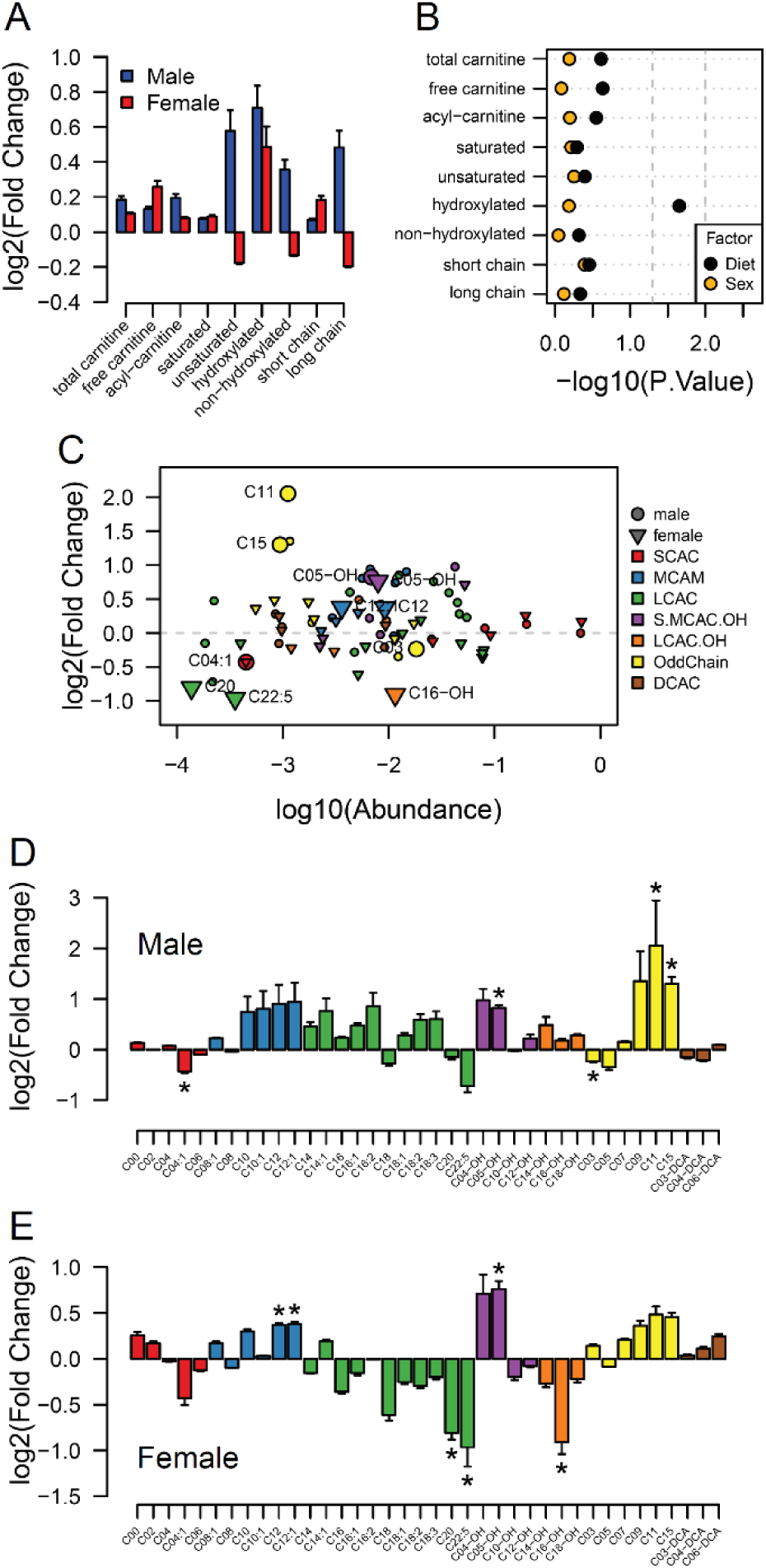
Acyl-carnitine levels in fetal hearts measured by LC-MS. (A) Relative changes in cardiac carnitine classes levels in male and female fetuses from obese dams. (B) Influence of maternal diet and sex on fetal cardiac carnitine classes levels as calculated by factorial ANOVA. (C) Relative fold change of individual acyl-carnitine levels in the heart of E18.5 fetuses from obese dams according to their abundance. Larger figures are acyl-carnitine species deemed as regulated with p<0.05 by Student t-test or Mann-Whitney test. SCAC: small-chain acyl-carnitine; MCAC: medium-chain acyl-carnitine; LCAC: long-chain acyl-carnitine; S.MCAM.OH: small- and medium-chain hydroxy acyl-carnitine, LCAC.OH: Long-chain hydroxy acyl-carnitine; Odd Chain: acyl-carnitines with an odd chain number; DCAC: dicarboxylic acylcarnitines. (D-E) Individual acyl-carnitine species levels in male (D) and female (E) fetal hearts at 18.5 days of pregnancy. See supplementary file 3 for list of full names. *p<0.05 by Student t-test or Mann-Whitney test. Male fetuses CTL n=7, male fetuses OB n=7, female fetuses CTL n=7, female fetuses OB n=6. Full data is available in Figure 6–source Data 1.

## DISCUSSION

In this study, we investigated the effect of an obesogenic *in utero* environment on the fetal cardiac and serum lipidome and explored if any effects were sexually dimorphic. We report for the first time unique patterns in the lipidome of the fetal heart as a consequence of maternal obesity which, despite exposure to the same *in utero* environment and similar serum lipid profiles, differed between male and female fetuses. The findings were consistent with changes in substrate availability that may affect fetal cardiac gene expression which is known to change in late gestation in preparation for birth.

Using direct infusion lipidomics, we observed unique responses to a maternal obesogenic diet between dams and offspring. This is consistent with the suggestion that the mouse placenta selectively transports lipids to the fetus, adjusting placental transfer to changes in maternal status (Miranda, 2018). In this process, maternal lipoproteins are hydrolysed at the placenta and non-esterified fatty acids are released into the fetal circulation bound to alpha-fetoproteins. Fatty acids are then taken up by the fetal liver and incorporated into lipoproteins, which are released into the fetal circulation (Reviewed by Herrera and Desoye, 2016). Thus, a combination of selective fatty acid transport by the placenta and fetal metabolism can explain the difference between maternal and fetal serum lipid profiles in obese pregnancies.

The analysis of both male and female fetal hearts and serum is a major novelty and strength of this study, allowing for investigation of any sex-specific differences in response to maternal obesity. There is growing evidence to suggest that there are sex-specific differences in response to a suboptimal *in utero* environment or at least in the timing of the development of the phenotype, with the male fetus being generally more vulnerable to the long-term detrimental consequences (Nicholas *et al,* 2020; Dearden *et al,* 2018). In the current study, a greater number of significant differences between control and obese cardiac lipidomes were observed in female fetuses than were seen in males. This difference between the sexes is particularly striking, given that fetuses of both sexes are exposed to the same maternal metabolic milieu. It is not clear if these sexually dimorphic responses early in life could represent an ability of females to adapt to the environment and to protect against longer term detrimental effects. Furthermore, although several individual lipid species displayed sex-dependent regulation, a similar overall impact of maternal obesity was observed in the serum lipidomes of both male and female offspring. Therefore, there were tissue specific effects of maternal obesity on the fetal serum and cardiac lipidomes. In the serum, triglycerides were more abundant, and several phosphatidylcholines/odd chain phosphatidylethanolamines were less abundant in response to maternal obesity.

The observed increase in fetal serum triglyceride and decrease in phosphatidylcholine/odd chain phosphatidylethanolamines abundances is suggestive of a change in lipoprotein composition. Previous studies showed that hyperlipidaemic human serum samples have a differential increase in the concentration of lipid species, with triglycerides increasing several fold whereas phosphatidylcholine only increased modestly (Kuklenyik *et al*, 2018). Although lipoprotein monolayers consist primarily of phosphatidylcholines enclosing a hydrophobic core containing triglycerides and cholesterol (Reviewed by Van der Veen *et al*, 2017), we do not necessarily expect a positive correlation between these two lipid classes. Lipoproteins are not always spherical and thus the relationship between particle surface area and volume is not uniform. Furthermore, high density lipoproteins (HDLs) comprise mainly phosphatidylcholines and very little triglyceride and, as the primary lipoprotein particles produced by the fetal liver are HDLs (Herrera and Desoye, 2016), a decrease in phosphatidylcholines without a corresponding decrease in triglyceride abundance may indicate a relative decrease in fetal HDL levels. A relative increase in triglyceride abundance in the absence of a relative increase in phosphatidylcholine may also indicate an increase in lipoprotein particle size with a smaller surface/core volume ratio in obese offspring (Kuklenyik *et al*, 2018).

Regardless of the precise mechanisms involved, the fetal serum signature indicates that maternal obesity causes deep changes to the lipids in the circulation of fetuses and thus those available to be taken up by fetal tissues. We observed that DPA – a 22:5 long-chain polyunsaturated fatty acid – from phospholipids was consistently lower in obese dams and in the serum and cardiac tissues of fetuses of both sexes. DPA is highly abundant in fish oils but can also be synthesized endogenously through eicosapentaenoic acid and arachidonic acid metabolism (Burdge *et al.,* 2002). Lower serum levels of this fatty acid have recently been associated with increased markers of insulin resistance and with increased cardiovascular risk in pregnant women (Zhu *et al*, 2019). However, as far as we are aware, a direct relationship between this fatty acid and fetal health has not yet been drawn, and we believe we are the first to identify a relative reduction of phospholipid-derived DPA levels in multiple fetal compartments. Moreover, similar changes in serum and cardiac fatty acid composition of lipids might also indicate that the fetal myocardium exposed to maternal-obesity-induced stress maintains its ability to uptake circulating fatty acids.

We observed changes in the fetal cardiac transcriptome that were induced by maternal obesity, including changes in regulation of genes associated with sterol, fatty acid and carnitine metabolism, that would indicate an early shift towards fatty acid oxidation. The fetal heart relies predominantly on glycolytic metabolism, however, at birth there is a switch to lipid oxidative metabolism. Increasing oxygen levels and high lipid availability in the maternal milk both play a crucial role in activating pathways that will ultimately lead to metabolic, physiological and morphological changes, resulting in postnatal heart maturation (Piquereau and Ventura-Clapier, 2018). The current observations therefore suggest that fatty acid availability drives a premature switch in metabolism from glucose to fatty acid oxidation in the fetal heart through PPAR-alpha activation by ligands, such as docosahexaenoic acid (22:6). This indicates early metabolic, though not morphological, maturation due to a change in the availability of nutrients to fetal tissues from obese pregnancies, which could have a long-term impact on cardiac function.

Previous studies have suggested that maternal obesity results in placental hypoxia and a reduction in the availability of oxygen to other fetal tissues (Wallace *et al*, 2019). Consistently, we previously observed increased HIF1A protein in the obese placenta (Fernandez-Twinn *et al*., 2017), indicating lower oxygen diffusion to the fetal tissues which would be expected to impair fatty acid oxidation. Indeed, despite the increased expression of *Cpt* genes, we observed cardiac accumulation of total hydroxy acyl-carnitine, an intermediate of beta-oxidation (Ventura *et al*, 1998). CPT proteins are required for fatty acid mitochondrial import and oxidation through addition and removal of carnitine from acyl groups, allowing their transport through the intermembrane space. Accumulation of total hydroxy acyl-carnitine is also observed in diabetic hearts (Su *et al*, 2005), and may indicate impaired mitochondrial capacity to completely metabolize the fatty-acid surplus in the matrix. Therefore, the expression of lipolytic genes alone may not be sufficient to compensate for the surplus in fatty acids and to increase the energy production and contractile potential of the fetal heart in a hypoxic environment. This may contribute to the reduced cardiomyocyte cell size, as observed in the female hearts from obese mothers. Nevertheless, further studies are required to define the fatty acid oxidation rates and energy balance in the fetal heart.

In conclusion, we have carried out a comprehensive study of how obesity during pregnancy influences lipid availability to the fetus and consequently affects the fetal heart lipidome. From our findings, three main principles emerge: (1) There is a discrepancy between how the maternal metabolic status affects the maternal and fetal serum lipidome, an outcome likely related both to the placental actions as a selective barrier and to changes in fetal metabolism; (2) Despite being exposed to the same maternal metabolic milieu, male and female fetal hearts show distinct responses to maternal obesity, mainly at the level of fatty acid residues and individual lipid isoforms. Female heart lipidomes were generally more sensitive and exhibited greater changes than males. Cardiac morphological changes were also uniquely observed in females at this age; (3) Changes in lipid supply resulting from maternal obesity might contribute to early expression of lipolytic genes in mouse hearts, possibly contributing to the previously observed changes in heart function in adult life. The precise mechanisms by which these alterations impact on long term cardiovascular health across the life course remains to be determined.

## MATERIAL AND METHODS

### Lead contact

Further information and requests for resources and reagents should be directed to and will be fulfilled by the Lead Contact, Lucas Carminatti Pantaleão.

### Data and Code Availability

The transcriptomics and lipidomics datasets generated during this study are available at GEO [GSE162185] and as supplemental material (Figure 2–source data 1, Figure 3–source data 1, Figure 4–source data 1, Figure 4–source data 2 and Figure 6 – source data 1), respectively.

### Animal handling

This research was regulated under the Animals (Scientific Procedures) Act 1986 Amendment Regulations 2012 following ethical review by the University of Cambridge Animal Welfare and Ethical Review Body (AWERB). Female C57BL/6J mice were randomly allocated to receive either commercial standard (RM1) or a high fat diet [20% lipids (Special Dietary Services)] plus sweetened condensed milk [55% simple carbohydrates/8% lipids (Nestle, UK)] from weaning. After around 6 weeks on their respective diets, mice were mated with male counterparts and went through a first pregnancy and lactation to ensure their breeding competence. Mice were mated a second time, and first day of pregnancy was marked by the detection of a vaginal plug. On day 18.5 of gestation, pregnant mice were culled by rising CO_2_ concentrations or cervical dislocation. Fetuses were surgically removed and immediately euthanised by laying in cold buffer. Fetal heart ventricles were dissected, weighed, snap frozen and stored at −80 °C or whole fetal torsos were fixed in 10% neutral buffered formalin. Maternal blood was collected by cardiac puncture and fetal trunk blood was collected following *post-mortem* head removal. All blood samples were processed for serum separation following standard protocol.

### Histology

Fixed fetal torsos were processed, embedded in paraffin and sectioned in the coronal plane at 10µm using a microtome (Leica Microsystems). Two to three slides for each fetus containing mid-cardiac sections were selected, processed, and stained with haematoxylin and eosin. Sections were imaged using a Slide Scanner Axio Scan Z1 (Zeiss, Germany). Nuclei counting analyses were performed blinded using open source digital analysis software (QuPath v0.2.0; Bankhead et al., 2017). The total cardiac region was manually selected, with the epicardium excluded, and colour deconvolution applied to the image to optimise stain separation. A composite training image consisting of randomly sampled regions from a subset of images was used to train a classifier to identify cardiac tissue from lumen and blood vessels, and nuclei detection was run on the classified tissue using automatic cellular detection after parameter optimisation (setup parameters of detection image: haematoxylin OD, pixel size: 0.5µm; nucleus parameters of background radius: 7µm, median filter radius: 0µm, sigma: 2µm, minimum area: 10µm^2^, maximum area: 400µm^2^, threshold: 0.15, max background intensity: 2, split by shape: TRUE). Automatic cellular detection was validated for a subset of images by manual counting of nuclei for random fields of view which sampled approximately 5-10% of the tissue.

### qPCR and RNA Sequencing

Cardiac RNA from male fetuses was extracted using a Qiazol/miRNeasy mini kit protocol (Qiagen, Hilden, Germany). Library preparation for mRNA sequencing followed manufacturer protocol of TruSeq RNA Library Preparation kit (Illumina, Cambridge, UK). Libraries were sequenced using a HiSeq 4000 platform and raw reads were mapped to mouse genome through bowtie v1.2.3. For RT-qPCR, RNA was reverse-transcribed using a High-Capacity cDNA Reverse Transcription Kit (Thermo-Fisher, Waltham, MA, USA). QPCR reactions were prepared using SYBR Green Master Mix (Thermo-Fisher) and specific primers. Fold changes were calculated by the 2−ΔΔCT method using *Hprt* and *Sdha* as housekeeping genes. Primers and oligonucleotides used in this study are listed in Figure 5–source data 1.

### Untargeted Lipidomics – Preparation of Samples

Solvents were purchased from Sigma-Aldrich Ltd (Dorset, UK) of at least HPLC grade and were not purified further. Lipid standards were purchased from Avanti Polar lipids (Alabaster, AL; via Instruchemie, Delfzijl, NL) and C/D/N/ isotopes (Quebec, Canada; via Qmx Laboratories, Thaxted, UK) and used without purification. Consumables were purchased from Sarstedt AG & Co (Leicester, UK) or Wolf Labs (Wolverhampton, UK).

The methods for preparing samples and extracting lipids used was described recently (Furse *et al*., 2020). Briefly, frozen whole fetal hearts were homogenised in a stock solution of guanidine and thiourea (6M/1·5M; 20× *w/v*) using mechanical agitation. The dispersions were freeze-thawed once before being centrifuged (12,000 × *g*, 10 min). The supernatant was collected and frozen (−80 °C) immediately. The thawed dispersion (60 µL) and serum aliquots (20 µL) were injected into wells (96w plate, Esslab Plate+™, 2·4 mL/well, glass-coated) followed by internal standards (150 µL, mixture of Internal Standards in methanol), water (500 µL) and DMT (500 µL, dichloromethane, methanol and triethylammonium chloride, 3:1:0·005). The mixture was agitated (96 channel pipette) before being centrifuged (3·2 × g, 2 min). A portion of the organic solution (20 µL) was transferred to an analytical plate (96w, glass coated, Esslab Plate+™) before being dried under Nitrogen gas. The dried films were re-dissolved (TBME, 30 µL/well) and diluted with a stock mixture of alcohols and ammonium acetate (100 µL/well; propan-2-ol: methanol, 2:1; CH3COO.NH4, 7·5 mM). The analytical plate was heat-sealed and run immediately.

### Untargeted Lipidomics – Direct Infusion Mass Spectrometry

Samples were directly infused into an Exactive Orbitrap (Thermo, Hemel Hampstead, UK), using a TriVersa NanoMate (Advion, Ithaca, US) for Direct Infusion Mass Spectrometry (Harshfield *et* al., 2019, Furse *et al*., 2020). A three-part analytical method was used (Furse *et al*., 2020, Furse and Koulman *et* al., 2019) in which samples were ionised in positive, negative, and then negative-with collision-induced-ionisation modes. The NanoMate infusion mandrel was used to pierce the seal of each well before an aliquot of the solution (15 μL) was collected with an air gap (1·5 μL). The tip was pressed against a fresh nozzle and the sample was dispensed using 0·2 psi (N_2 (g)_). Ionisation was achieved at a 1·2 kV. The Exactive was set to start acquiring data 20 s after sample aspiration began. The data were collected with a scan rate of 1 Hz (resulting in a mass resolution of 65,000 full width at half-maximum (fwhm) at 400 *m/z*). After 72 s of acquisition in positive mode the NanoMate and the Exactive switched to negative mode, decreasing the voltage to −1·5 kV. The spray was maintained for another 66 s, after which Collision-Induced Dissociation (CID) commenced, with a mass window of 50–1000 Da, and was stopped after another 66 s. The analysis was then stopped, and the tip discarded before the analysis of the next sample began. The sample plate was kept at 10 °C throughout the data acquisition. Samples were run in row order.

Raw high-resolution mass-spectrometry data were processed using XCMS (www.bioconductor.org) and Peakpicker v 2.0 (an in-house R script, Harshfield *et* al., 2019). Theoretical lists of known species (by m/z) were used for both positive ion and negative ion mode (∼8·5k species including different adducts and fragmentations). Variables whose mass deviated by more than 9 ppm from the expected value, had a signal-to-noise ratio of <3 and had signals for fewer than 20% of samples were discarded. The correlation of signal intensity to concentration of lipid variables found in pooled mouse heart homogenate, pooled mouse liver homogenate, and pooled human serum samples (0·25, 0·5, 1·0×) was used to identify the lipid signals the strength of which was linearly proportional to abundance (r > 0·75) in samples.

For the detection of fatty acids of phospholipids only, a deviations threshold of 12·5ppm was used for processing of the negative mode with CID, on a list of fatty acids of chain length 14 to 36 with up to six double bonds and/or one hydroxyl group. All signals greater than noise were carried forward. In this method the lipidome is not separated chromatographically but measured only by mass-to-charge ratio and therefore cannot distinguish lipids that are isobaric (identical molecular mass) in a given ionisation mode. In this study, the identification of the lipids was based on their accurate mass in positive ionization mode according to Lipid Maps structure database (Sud *et al*., 2007). In case of multiple isobars per lipid signal, the likely identification was predicted according to the biological likelihood, full list of possible annotations can be found in the supplementary information (Supplementary file 1 and supplementary file 2). Signals consistent with fatty acids were found in 3/3 samples checked manually. Relative abundance was calculated by dividing each signal by the sum of signals for that sample, expressed per mille (‰). Zero values were interpreted as not measured and for the remaining non-assigned values, we used a known single component projection based on nonlinear iterative partial least squares algorithm (Nelson *et al*., 1996) to impute values and populate the dataset. Data was normalized using quantile Cyclic Loess method and statistical calculations were done on these finalised values.

### Acyl-carnitine Analysis – Preparation of Samples

All solvents and additives were of HPLC grade or higher and purchased from Sigma Aldrich unless otherwise stated.

The protein precipitation liquid extraction protocol was as follows: the tissue samples were weighed (between 1.4 - 11.0 mg) and transferred into a 2 mL screw cap Eppendorf plastic tubes (Eppendorf, Stevenage, UK) along with a single 5 mm stainless steel ball bearing. Immediately, 400 µL of chloroform and methanol (2:1, respectively) solution was added to each sample, followed by thorough mixing. The samples were then homogenised in the chloroform and methanol (2:1, respectively) using a Bioprep 24-1004 homogenizer (Allsheng, Hangzhou City, China) run at speed; 4.5 m/s, time; 30 seconds for 2 cycles. Then, 400 µL of chloroform, 100 µL of methanol and 100 µL of the stable isotope labelled acyl-carnitine internal standard; containing butyryl-d7-L-carnitine (order number: D-7761, QMX Laboratories Ltd. (QMX Laboratories Ltd, Essex, United Kingdom)) and hexadecanoylLcarnitine-d3 (order number: D-6646, QMX Laboratories Ltd.) at 5 µM in methanol was added to each sample. The samples were homogenised again using a Bioprep 24-1004 homogenizer run at speed; 4.5 m/s, time; 30 seconds for 2 cycles. To ensure fibrous material was diminished, the samples were sonicated for 30 minutes in a water bath sonicator at room temperature (Advantage-Lab, Menen, Belgium). Then, 400 µL of acetone was added to each sample. The samples were thoroughly mixed and centrifuged for 10 minutes at ∼20,000 g to pellet any insoluble material at the bottom of the vial. The single layer supernatant was pipetted into separate 2 mL screw cap amber-glass autosampler vials (Agilent Technologies, Cheadle, United Kingdom); being careful not to break up the solid pellet at the bottom of the tube. The organic extracts (chloroform, methanol, acetone composition; ∼7:3:4, ∼1.4 mL) were dried down to dryness using a Concentrator Plus system (Eppendorf, Stevenage, UK) run for 60 minutes at 60 degree Celsius. The samples were reconstituted in 100 µL of water and acetonitrile (95:5, respectively) then thoroughly mixed. The reconstituted sample was transferred into a 250 μL low-volume vial insert inside a 2 mL amber glass auto-sample vial ready for liquid chromatography with mass spectrometry detection (LC-MS) analysis.

### Acyl-carnitine Analysis – Liquid Chromatography Mass Spectrometry

Full chromatographic separation of acyl-carnitines was achieved using Shimadzu HPLC System (Shimadzu UK Ltd., Milton Keynes, United Kingdom) with the injection of 10 µL onto a Hichrom ACE Excel 2 C18-PFP column (Hichrom Ltd., Berkshire, United Kingdom); 2.0 µm, I.D. 2.1 mm X 150 mm, maintained at 55 °C. Mobile phase A was water with 0.1% formic acid. Mobile phase B was acetonitrile with 0.1% formic acid. The flow was maintained at 500 µL per minute through the following gradient: 0 minutes_5% mobile phase B, at 0.5 minutes_100% mobile phase B, at 5.5 minutes_100% mobile phase B, at 5.51 minutes_5% mobiles phase B, at 7 minutes_5% mobile phase B. The sample injection needle was washed using acetonitrile and water mix (1:1, respectively). The mass spectrometer used was the Thermo Scientific Exactive Orbitrap with a heated electrospray ionisation source (Thermo Fisher Scientific, Hemel Hempstead, UK). The mass spectrometer was calibrated immediately before sample analysis using positive and negative ionisation calibration solution (recommended by Thermo Scientific). Additionally, the heated electrospray ionisation source was optimised to ensure the best sensitivity and spray stability (capillary temperature; 300 degree Celsius, source heater temperature; 420 degree Celsius, sheath gas flow; 40 (arbitrary), auxiliary gas flow; 15 (arbitrary), spare gas; 3 (arbitrary), source voltage; 4 kV. The mass spectrometer scan rate set at 2 Hz, giving a resolution of 50,000 (at 200 m/z) with a full-scan range of m/z 150 to 800 in positive mode.

Thermo Xcalibur Quan Browser data processing involved the integration of the internal standard extracted ion chromatogram (EIC) peaks at the expected retention times: butyryl-d7-L-carnitine ([M+H]+, m/z 239.19827 at 1.20 minutes) and hexadecanoyl-L-carnitine-d3 ([M+H]+, m/z 403.36097 at 4.20 minutes). The data processing also involved the integration of the targeted individual acyl-carnitine species (m/z was [M+H]+) at their expected retention time allowing for a maximum of ±0.1 minutes of retention time drift: any retention time drift greater than ±0.1 minutes resulted in the exclusion of the analyte leading to a ‘Not Found’ result (i.e., zero concentration). Through the Thermo Xcalibur Quan Browser software the responses of the analytes were normalised to the relevant internal standard response (producing area ratios), these area ratios corrected the intensity for any extraction and instrument variations. The area ratios were then blank corrected where intensities less than three times the blank samples were set to a ‘Not Found’ result (i.e., zero concentration). The accepted area ratios were then multiplied by the concentration of the internal standard (5 µM) to give the analyte concentrations. For tissue samples, the calculated concentrations (µM) of the analytes were then divided by the amount of tissue (in mg) used in the extraction protocol to give the final results in µM per mg of tissue extracted (µM/mg).

### Sample-size estimation

Due to the untargeted high-throughput aspect of this study and to the scarcity of available data into fetal lipidomics, the use of a power analysis accounting for changes in fetal cardiac lipids to predict sample size was challenging. The number of animals used in the present study was therefore predicted using previously obtained data on histological assessment of the ratio left ventricle:lumen in the male fetal heart, and on the extensive track record of published studies from our research group using the same maternal obesity model employed in the current study. According to an *a priori* unpaired t-test power calculation, an n equal or greater than 5 would be required to achieve significance set at α < 0.05, 80% power. Also, according to the resource equation, an n equal to or greater than 6 results in more than 10 degrees of freedom, and is therefore adequate. We then concluded that a sample size greater than 6 would be necessary to show any significant changes in our study.

### Biometric markers and qPCR – Statistical Analysis

Details of statistical analysis (statistical tests used, number of animals and precision measures) can be found in the figure legends. Simple Student t-test was employed to identify statistically significant differences in all biometric, histological and qPCR analysis, comparing control and obese groups in a sex-dependent manner. Factorial ANOVA was employed to test offspring sex and maternal status influence on individual mRNA levels.

### RNASeq – Statistical Analysis

Reads per Kilobase of transcript per Million mapped reads (RPKM) were produced from RNA Sequencing raw output and statistically analysed through likelihood ratio test using R version 3.6.3. Core analysis in Ingenuity® Pathway Analysis application (IPA – Qiagen) was used in data interpretation and pathway enrichment. A p-value cut-off of 0.01 was used to determine genes to be mapped to IPA networks.

### Untargeted and Targeted Lipidomics – Statistical Analysis

Uni- and multivariate statistical models were created using R version 3.6.3. Multiple Shapiro-Wilk tests were carried out to identify if individual variables were normally distributed. Multiple t-tests were used to identify significant regulation of individual lipid species, and multiple t-tests or Mann-Whitney tests were used to identify individual lipid classes differences between groups when individual variables were normally or non-normally distributed. A multivariate partial-least square discriminatory analysis (PLS-DA) was also employed to identify Variable Importance in the Projection (VIP) and determine individual lipids that maximise the model classification ability. For individual lipid species, variables were deemed significantly regulated and relevant when p-value < 0.05 and vip score > 1. Individual lipid classes, acyl-carnitines and fatty acids were significantly regulated when p < 0.05. Factorial ANOVA was also used to test offspring sex and maternal status influence over individual lipid classes, and pools of fatty acids and acyl-carnitines. Prior to statistical analysis, outlier samples were identified through a combination of frequency distribution analysis, lipid classes frequency investigation, PCA and hierarchical clustering analysis. Samples with lower than 66.7% of lipid signals detected or deemed as outliers in all the aforementioned analyses failed the quality control for mass spectrometry and were excluded from the datasets and from further statistical tests. Lipid ontology enrichment analysis was carried out using LION (Molenaar *et al*., 2019). Lipid traffic analysis was conducted following previously described methods (Furse *et al*., 2020).

## COMPETING INTERESTS

The authors declare no competing interests.

## SUPPLEMENTAL MATERIAL LEGENDS

Supplementary file 1. Isobars and main predicted classes for m/z detected in direct infusion high-resolution mass spectrometry of the serum (positive mode only). Isobar annotations were obtained from LIPID MAPS Structure Database and a mass tolerance (m/z) threshold: +/− 0.001 was used. For multiple isobars per m/z, biological likelihood was employed to predict the likely identification. Main classes were predicted according to the likely identification.

Supplementary file 2. Isobars and main predicted classes for m/z detected in direct infusion high-resolution mass spectrometry of the heart (positive mode only). Isobar annotations were obtained from LIPID MAPS Structure Database and a mass tolerance (m/z) threshold: +/− 0.001 was used. For multiple isobars per m/z, biological likelihood was employed to predict the likely identification. Main classes were predicted according to the likely identification.

Supplementary file 3. List of names for acyl-carnitines identified in E18.5 fetal hearts by LCMS

Figure 2–source data 1. Direct infusion high-resolution mass spectrometry of the serum (positive mode only). Scaled raw data and statistical significance.

Figure 3–source data 1. Fatty acids abundance obtained by direct infusion high-resolution mass spectrometry of the serum (negative mode). Scaled raw data and statistical significance.

Figure 4–source data 1. Direct infusion high-resolution mass spectrometry of the heart (positive mode only). Scaled raw data and statistical significance.

Figure 4–source data 2. Fatty acids abundance obtained by direct infusion high-resolution mass spectrometry of the heart (negative mode). Scaled raw data and statistical significance.

Figure 5–source data 1. Sequence-specific primers for qPCR.

Figure 6–source data 1. Acyl-Carnitines abundance obtained by spectrometry of the heart (negative mode). Raw data and statistical significance.

## Notes

### Competing Interest Statement

The authors have declared no competing interest.

## REFERENCES

Bankhead, P., Loughrey, M.B., Fernández, J.A., Dombrowski, Y., McArt, D.G., Dunne, P.D., McQuaid, S., Gray, R.T., Murray, L.J., Coleman, H.G., James, J.A., Salto-Tellez, M., and Hamilton, P.W. (2017). QuPath: Open source software for digital pathology image analysis. Sci. Rep. 7(1), 16878. doi: 10.1038/s41598-017-17204-5.

Barker, D.J. (2007). The origins of the developmental origins theory. J. Intern. Med 261(5), 412–17. doi: 10.1111/j.1365-2796.2007.01809.x.

Burdge, G., Jones, A., and Wootton, S. (2002). Eicosapentaenoic and docosapentaenoic acids are the principal products of α-linolenic acid metabolism in young men. Br. J. Nutr. 88(4), 355–363. doi: 10.1079/BJN2002662.

Catalano, P. M., and Shankar, K. (2017). Obesity and pregnancy: mechanisms of short term and long term adverse consequences for mother and child. BMJ (Clinical research ed.), 356. doi: 10.1136/bmj.j1.

Dearden, L., Bouret, S. G., and Ozanne, S. E. (2018). Sex and gender differences in developmental programming of metabolism. Mol. Metab. 15, 8–19. doi: 10.1016/j.molmet.2018.04.007.

Dong, M., Zheng, Q., Ford, S.P., Nathanielsz, P.W., Ren, J. (2012). Maternal obesity, lipotoxicity and cardiovascular diseases in offspring. J. Mol. Cell Cardiol. 55, 111–6. doi: 10.1016/j.yjmcc.2012.08.023.

Eum, J., Lee, J., Yi, S., Kim, I., Seong, J. and Moon, M. (2020). Aging-related lipidomic changes in mouse serum, kidney, and heart by nanoflow ultrahigh-performance liquid chromatography-tandem mass spectrometry. J. Chromatogr. 1618, 460849. doi: 10.1016/j.chroma.2020.460849.

Fernandez-Twinn, D.S., Gascoin, G., Musial, B., Carr, S., Duque-Guimaraes, D., Blackmore, H.L., Alfaradhi, M.Z., Loche, E., Sferruzzi-Perri, A.N., Fowden, A.L., Ozanne, S.E. (2017). Exercise rescues obese mothers’ insulin sensitivity, placental hypoxia and male offspring insulin sensitivity. Sci. Rep. 7, 44650. doi: 10.1038/srep44650.

Furse, S., Fernandez-Twinn, D.S., Jenkins, B., Meek, C.L., Williams, H.E.L., Smith, G.C.S., Charnock-Jones, D.S., Ozanne, S.E., Koulman, A. (2020). A high-throughput platform for detailed lipidomic analysis of a range of mouse and human tissues. Anal. Bioanal. Chem. 412(12), 2851–2862. doi: 10.1007/s00216-020-02511-0.

Furse, S., Koulman, A. (2019). The Lipid and Glyceride Profiles of Infant Formula Differ by Manufacturer, Region and Date Sold. Nutrients 11(5), 1122. doi: 10.3390/nu11051122.

Furse, S., Watkins, A., Hojat, N., Smith, J., Williams, H., Chiarugi, D., & Koulman, A. (2021). Lipid traffic analysis reveals the impact of high paternal carbohydrate intake on offsprings’ lipid metabolism. Comms Biol., 4, 163. doi: 10.1038/s42003-021-01686-1.

GBD 2017 Causes of Death Collaborators (2018). Global, Regional, and National Age-Sex-Specific Mortality for 282 Causes of Death in 195 Countries and Territories, 1980-2017: A Systematic Analysis for the Global Burden of Disease Study 2017. Lancet 392(10159), 1736–1788. doi: 10.1016/S0140-6736(18)32203-7.

Guénard, F., Deshaies, Y., Cianflone, K., Kral, J.G., Marceau, P., Vohl, M.C (2013). Differential methylation in glucoregulatory genes of offspring born before vs. after maternal gastrointestinal bypass surgery. Proc. Natl. Acad. Sci 110(28), 11439–44. doi: 10.1073/pnas.1216959110.

Halade, G., Dorbane, A., Ingle, K., Kain, V., Schmitter, J. and Rhourri-Frih, B. (2018). Comprehensive targeted and non-targeted lipidomics analyses in failing and non-failing heart. Anal Bioanal Chem 410(7), 1965–1976. doi: 10.1007/s00216-018-0863-7.

Hamilton, R. and Fielding, P. (1989). Nascent very low density lipoproteins from rat hepatocytic Golgi fractions are enriched in phosphatidylethanolamine. Biochem. Biophys. Res. Comm 160(1), 162–167. 10.1016/0006-291x(89)91635-5.

Harshfield, E.L., Koulman, A., Ziemek, D., Marney, L., Fauman, E.B., Paul, D.S., Stacey, D., Rasheed, A., Lee, J.J., Shah, N., Jabeen, S., Imran, A., Abbas, S., Hina, Z., Qamar, N., Mallick, N.H., Yaqoob, Z., Saghir, T., Rizvi, S.N.H., Memon, A., Rasheed, S.Z., Memon, F.U., Qureshi, I.H., Ishaq, M., Frossard, P., Danesh, J., Saleheen, D., Butterworth, A.S., Wood, A.M., Griffin, J.L. (2019). An Unbiased Lipid Phenotyping Approach To Study the Genetic Determinants of Lipids and Their Association with Coronary Heart Disease Risk Factors. J. Proteome Res. 18(6), 2397–2410. doi: 10.1021/acs.jproteome.8b00786.

Helle, E. and Priest, J. (2020). Maternal Obesity and Diabetes Mellitus as Risk Factors for Congenital Heart Disease in the Offspring. J. Am. Heart Assoc. 9(8). doi: 10.1161/JAHA.119.011541.

Herrera, E., Desoye, G. (2016). Maternal and fetal lipid metabolism under normal and gestational diabetic conditions. Horm. Mol. Biol. Clin. Investig. 26(2), 109–27. doi: 10.1515/hmbci-2015-0025.

Howell, K. R. and Powell, T. L. (2017). Effects of maternal obesity on placental function and fetal development. Reproduction (Cambridge, England), 153(3), R97–R108. doi: 10.1530/REP-16-0495.

Kuklenyik, Z., Jones, J., Gardner, M., Schieltz, D., Parks, B., Toth, C., Rees, J., Andrews, M., Carter, K., Lehtikoski, A., McWilliams, L., Williamson, Y., Bierbaum, K., Pirkle, J. and Barr, J. (2018). Core lipid, surface lipid and apolipoprotein composition analysis of lipoprotein particles as a function of particle size in one workflow integrating asymmetric flow field-flow fractionation and liquid chromatography-tandem mass spectrometry. PLOS ONE 13(4), e0194797. doi: 10.1371/journal.pone.0194797. eCollection 2018.

Le, C.H., Mulligan, C.M., Routh, M.A., Bouma, G.J., Frye, M.A., Jeckel, K.M., Sparagna, G.C., Lynch, J.M., Moore, R.L., McCune, S.A., et al. (2014). Delta-6-desaturase links polyunsaturated fatty acid metabolism with phospholipid remodeling and disease progression in heart failure. Circ. Heart Fail. 7, 172–183. doi: 10.1161/CIRCHEARTFAILURE.113.000744.

Loche, E., Blackmore, H., Carpenter, A., Beeson, J., Pinnock, A., Ashmore, T., Aiken, C., de Almeida-Faria, J., Schoonejans, J., Giussani, D., Fernandez-Twinn, D. and Ozanne, S. (2018). Maternal diet-induced obesity programmes cardiac dysfunction in male mice independently of post-weaning diet. Cardiovasc. Res. 114(10), 1372–1384. doi: 10.1093/cvr/cvy082.

Lopaschuk, G. and Jaswal, J. (2010). Energy Metabolic Phenotype of the Cardiomyocyte During Development, Differentiation, and Postnatal Maturation. J. Cardiovasc. Pharmacol. 56(2), 130–140. doi: 10.1097/FJC.0b013e3181e74a14.

Miranda, J., Simões, R., Paules, C., Cañueto, D., Pardo-Cea, M., García-Martín, M., Crovetto, F., Fuertes-Martin, R., Domenech, M., Gómez-Roig, M., Eixarch, E., Estruch, R., Hansson, S., Amigó, N., Cañellas, N., Crispi, F. and Gratacós, E. (2018). Metabolic profiling and targeted lipidomics reveals a disturbed lipid profile in mothers and fetuses with intrauterine growth restriction. Sci. Rep. 8(1), 13614. doi: 10.1038/s41598-018-31832-5.

Molenaar, M.R., Jeucken, A., Wassenaar, T.A., van de Lest, C.H.A., Brouwers, J.F., Helms, J.B. (2019). LION/web: a web-based ontology enrichment tool for lipidomic data analysis. Gigascience 8(6), giz061. doi: 10.1093/gigascience/giz061.

Nelson, P.R.C., Taylor, P.A., MacGregor, J.F. (1996). Missing data methods in PCA and PLS: Score calculations with incomplete observations. Chemometr Intell Lab Syst. 35(1), 45–65. DOI:10.1016/S0169-7439(96)00007-X.

Nicholas, L. M., Nagao, M., Kusinski, L. C., Fernandez-Twinn, D. S., Eliasson, L., & Ozanne, S. E. (2020). Exposure to maternal obesity programs sex differences in pancreatic islets of the offspring in mice. Diabetologia, 63(2), 324–337. doi: 10.1007/s00125-019-05037-y.

NMPA Project Team (2019). National Maternity and Perinatal Audit: Clinical Report 2019. Based on births in NHS maternity services between 1 April 2016 and 31 March 2017 (London: RCOG).

Patterson, A.J., and Zhang, L. (2010). Hypoxia and fetal heart development. Curr. Mol. Med 10, 653–666.

Piquereau, J. and Ventura-Clapier, R. (2018). Maturation of Cardiac Energy Metabolism During Perinatal Development. Front. Physiol 9.

Su, X., Han, X., Mancuso, D., Abendschein, D. and Gross, R. (2005). Accumulation of Long-Chain Acylcarnitine and 3-Hydroxy Acylcarnitine Molecular Species in Diabetic Myocardium: Identification of Alterations in Mitochondrial Fatty Acid Processing in Diabetic Myocardium by Shotgun Lipidomics. Biochemistry 44(13), 5234–5245. doi: 10.1021/bi047773a.

Sud, M., Fahy, E., Cotter, D., Brown, A., Dennis, E.A., Glass, C.K., Merrill, A.H. Jr, Murphy, R.C., Raetz, C.R., Russell, D.W., Subramaniam, S. (2007). LMSD: LIPID MAPS structure database. Nucleic Acids Res. 35, D527–32. doi: 10.1093/nar/gkl838.

Tham, Y., Bernardo, B., Huynh, K., Ooi, J., Gao, X., Kiriazis, H., Giles, C., Meikle, P. and McMullen, J. (2018). Lipidomic Profiles of the Heart and Circulation in Response to Exercise versus Cardiac Pathology: A Resource of Potential Biomarkers and Drug Targets. Cell Rep. 24(10), 2757–2772.

van der Veen, J., Kennelly, J., Wan, S., Vance, J., Vance, D. and Jacobs, R. (2017). The critical role of phosphatidylcholine and phosphatidylethanolamine metabolism in health and disease. Biochim. Biophys. Acta 1859(9), 1558–1572. doi: 10.1016/j.celrep.2018.08.017.

Ventura, F., Ijlst, L., Ruiter, J., Ofman, R., Costa, C., Jakobs, C., Duran, M., De Almeida, I., Bieber, L. and Wanders, R. (1998). Carnitine palmitoyltransferase II specificity towards beta-oxidation intermediates. Evidence for a reverse carnitine cycle in mitochondria. Eur. J. Biochem 253(3), 614–618. doi: 10.1046/j.1432-1327.1998.2530614.x.

Wallace, J., Bellissimo, C., Yeo, E., Fei Xia, Y., Petrik, J., Surette, M., Bowdish, D. and Sloboda, D. (2019). Obesity during pregnancy results in maternal intestinal inflammation, placental hypoxia, and alters fetal glucose metabolism at mid-gestation. Sci. Rep. 9(1). doi: 10.1038/s41598-019-54098-x.

Zambrano, E., Nathanielsz, P.W. (2013). Mechanisms by which maternal obesity programs offspring for obesity: evidence from animal studies. Nutr. Rev. Suppl 1, S42–54. doi: 10.1111/nure.12068.

Zhu, Y., Li, M., Rahman, M., Hinkle, S., Wu, J., Weir, N., Lin, Y., Yang, H., Tsai, M., Ferrara, A. and Zhang, C. (2019). Plasma phospholipid n-3 and n-6 polyunsaturated fatty acids in relation to cardiometabolic markers and gestational diabetes: A longitudinal study within the prospective. PloS Med. 16(9), e1002910. doi: 10.1371/journal.pmed.1002910.

